# Direct serological antibody discovery by integrative proteomics yields potent neutralizers overlooked by single-cell BCR sequencing

**DOI:** 10.64898/2026.05.20.725438

**Authors:** Sem Tamara, Danique M.H. van Rijswijck, Jacqueline van Rijswijk, Judith A. Burger, Douwe Schulte, Marta Moreschini, Godelieve J. de Bree, Marit J. van Gils, Albert J.R. Heck, Maurits A. den Boer

**Affiliations:** Abvion Biologics B.V., Padualaan 8, 3584 CH, Utrecht, The Netherlands; Biomolecular Mass Spectrometry and Proteomics, Bijvoet Center for Biomolecular Research and Utrecht Institute for Pharmaceutical Sciences, Utrecht University, Padualaan 8, 3584 CH, Utrecht, The Netherlands; Netherlands Proteomics Center, Padualaan 8, 3584 CH, Utrecht, The Netherlands; Department of Medical Microbiology and Infection Prevention, Amsterdam UMC, University of Amsterdam, 1105 AZ, Amsterdam, The Netherlands; Amsterdam Institute for Immunology and Infectious Diseases, 1105 AZ, Amsterdam, The Netherlands; Department of Internal Medicine, Amsterdam UMC, University of Amsterdam, Amsterdam Institute for Infection and Immunity, 1105 AZ, Amsterdam, The Netherlands

## Abstract

Human antibody discovery relies on accessing *in vivo*-matured repertoires, yet conventional single-cell B cell receptor sequencing (scBCR-seq) often overlooks the most relevant, functional antibodies secreted by plasma cells. Here, we introduce AbDirect, a protein-centric discovery platform that obtains antibody sequences directly from small-volume biofluids. In this proof-of-concept, we apply AbDirect to potent COVID-19 plasma from which an early-pandemic scBCR-seq study did not identify neutralizers. Upfront reactivity screening of anti-SARS-CoV-2 spike protein repertoires revealed diverse clonal profiles with distinct cross-reactivity and subunit specificity. Targeted *de novo* sequencing via standalone integrative proteomics yielded 14 IgG1 and 4 IgA1 clones that diverged markedly from peripheral B cell counterparts in germline usage and phylogeny, indicating distinct immunological compartments. Validation via recombinant mAbs demonstrated superior binding and highly potent neutralization for multiple sequenced clones (three with IC_50_ ≤1.4 nM). AbDirect thus yielded potent antibodies overlooked by scBCR-seq, demonstrating serological discovery as a powerful complementary approach for uncovering functional repertoires that may be inaccessible to cell-based methods.

## Introduction

Monoclonal antibodies (mAbs) represent one of the most successful therapeutic formats, with five out of the ten best-selling drugs and a global market exceeding USD 280 billion in 2025^1–3^. Driven by these successes, direct access to *in vivo*-matured antibody repertoires has become a cornerstone of therapeutic development^4,5^. Natural human repertoires present sources of highly potent mAbs under wide-ranging conditions, including infectious diseases, autoimmune disorders, and cancer^6–8^. Discovery of such mAbs enables the development of therapies that broadly and potently neutralize pathogens, modulate dysregulated immune responses, or effectively target tumor cells^4,7^. For instance, in infectious diseases like COVID-19^9–12^, Ebola^13^, and RSV^14^, patient-derived antibodies have already been instrumental in reducing severe outcomes. Still, restricted access to relevant mature sequences limits therapeutic success and underscores the need for innovative platforms that efficiently capture functional repertoires from diverse humoral sources^15–17^.

The current state-of-the-art in antibody discovery does not directly assess serological antibodies but is instead dominated by single-cell B cell receptor sequencing (scBCR-seq)^5,18–21^. This powerful and widely successful approach captures BCR repertoires at the transcriptomic level and has enabled the identification of numerous potent therapeutic mAbs from convalescent donors. It excels at high-throughput sequencing of heavy and light chain pairs from antigen-specific memory B cells, facilitating the discovery of diverse antibodies. In humans, this is practically limited to peripheral blood mononuclear cells (PBMCs) from large blood draws, capturing primarily circulating memory B cells. These are, however, not the antibody-secreting long-lived plasma cells responsible for systemic antibodies present in serum, as the latter reside mostly in the bone marrow^18,22–24^. This indirect BCR-based strategy, requiring *in vitro* antibody reconstruction, may therefore not necessarily reflect the functional serological antibody repertoire^5,19,25^. Additional challenges include the potential inclusion of low-affinity or non-secreted sequences, the inability to assess clonal abundance, and incomplete capture of isotype diversity (e.g., IgA). Furthermore, resulting sequences often need extensive downstream validation and optimization, particularly in complex diseases such as autoimmunity and oncology, where appropriate peripheral BCRs can be challenging to find^26–30^.

Protein-based antibody discovery offers a compelling alternative strategy for identifying therapeutically relevant antibodies directly from circulation^31–37^. This approach identifies *in vivo*-matured, functional molecules directly through liquid chromatography-mass spectrometry (LC-MS) or cryogenic electron microscopy polyclonal epitope mapping (cryo-EMPEM) analysis of secreted antibodies. Through affinity enrichment of antibodies from their native, polyclonal context, this may preferentially yield high-relevance sequences with a reduced need for costly steps like *in vitro* affinity maturation and lowered risks of poor expression or stability^17,38^. The approach enables direct access to small-volume biofluids (e.g., 0.01-2 mL), bypassing the need for viable cells and making it suitable for retrospective studies and scenarios with limited fresh sampling material^39^. Furthermore, it preserves isotype diversity, PTM profiles, and, particularly by LC-MS, the ability for quantitative profiling of clonal dominance and assessing temporal dynamics^40–43^. Combined, this enables capturing a high-fidelity picture of genuine, functional serological antibody repertoires at the protein level, allowing the identification of potent candidates from human elite responders across disease areas^17,44,45^.

A major restriction to mass spectrometry-based analysis of serological antibody repertoires has been the critical challenge of obtaining individual antibody sequences from highly complex polyclonal mixtures. Yet, several recent advances in LC-MS technologies highlight the potential for protein-based antibody discovery^31,32,34,35,46^. Foundational work focused on bottom-up proteomics of monoclonal or fractionated antibodies, involving multiple protease digestion schemes and assembly of *de novo* sequenced peptides. In hybrid proteogenomics approaches, this is often combined with bulk or single-cell B cell sequencing data, with pioneering work already demonstrating the limited overlap between serum CDRH3 peptides and peripheral BCR sequences^17,31,47^. In recent advances, this approach has furthermore enabled the identification of up to three fully protein-derived antibody sequences from high-titer human plasma^34,35^. As an alternative standalone strategy, integrated proteomics may tackle sensitivity and coverage by layering peptide-level data with direct LC-MS measurements of intact antibodies and released chains^32,43,46,48,49^. This includes innovations for profiling serological repertoires as well as sequence discovery from endogenous samples^32,40,46,50^. Combined, these advances have laid a strong foundation, yet standalone *de novo* sequencing of dozens of full-length, chain-paired antibodies directly from polyclonal serum has remained elusive.

Here, we introduce a protein-centric antibody discovery platform (AbDirect), extending integrative proteomics into a standalone approach that resolves antibody sequence panels directly from complex polyclonal samples. In this proof-of-concept study, we apply and benchmark the platform on COVID-19 plasma from an early pandemic study where, despite robust serum activity, scBCR-seq did not yield any neutralizing antibodies^51^. AbDirect is positioned to address this specific scenario, exemplified here by donor COSCA3. From the exact same sample, we first screen the serological repertoire for clonal-level reactivities against SARS-CoV-2 spike (S) protein variants and subunits (i.e. RBD, S1, and S2), after which we fully sequence a polyclonal panel of 18 affinity-enriched antibodies. We next compare this serological panel to previous BCR-based outcomes, highlighting that this direct protein-centric approach delivers distinct sequences, including multiple high-affinity and potently neutralizing clones. AbDirect can be applied for retrieving antibodies using a wide range of biofluids and immobilized antigens. With current sensitivity enabling sequencing from as little as 1 µg of enriched material, future applications may include antibody discovery from archived biofluids across disease areas, supporting both repertoire analyses and therapeutic antibody development.

## Results

### Sample description and longitudinal repertoire profiling

For benchmarking the AbDirect approach *versus* conventional scBCR-seq, we selected plasma from donor COSCA3 from the study of Brouwer et al.^51^ (**Figure 1**). This 69-year-old male was severely affected by SARS-CoV-2 infection early in the pandemic (WT strain, March 2020), requiring ICU admission days 11-18 post symptom onset. At day 23, high serum titers and strong serum neutralization activity^a^ were observed^51^. Notably, while scBCR-seq on PBMCs from this exact sample yielded 105 unique VH and 27 unique VL sequences, recombinant expression and validation of the 10 pairs with the highest affinity for the S protein surprisingly revealed no neutralizing antibodies at all. We reasoned that this sample could serve as a compelling test case for a proof-of-concept study using protein-centric antibody discovery to recapitulate this robust serum activity.

**Figure 1.**
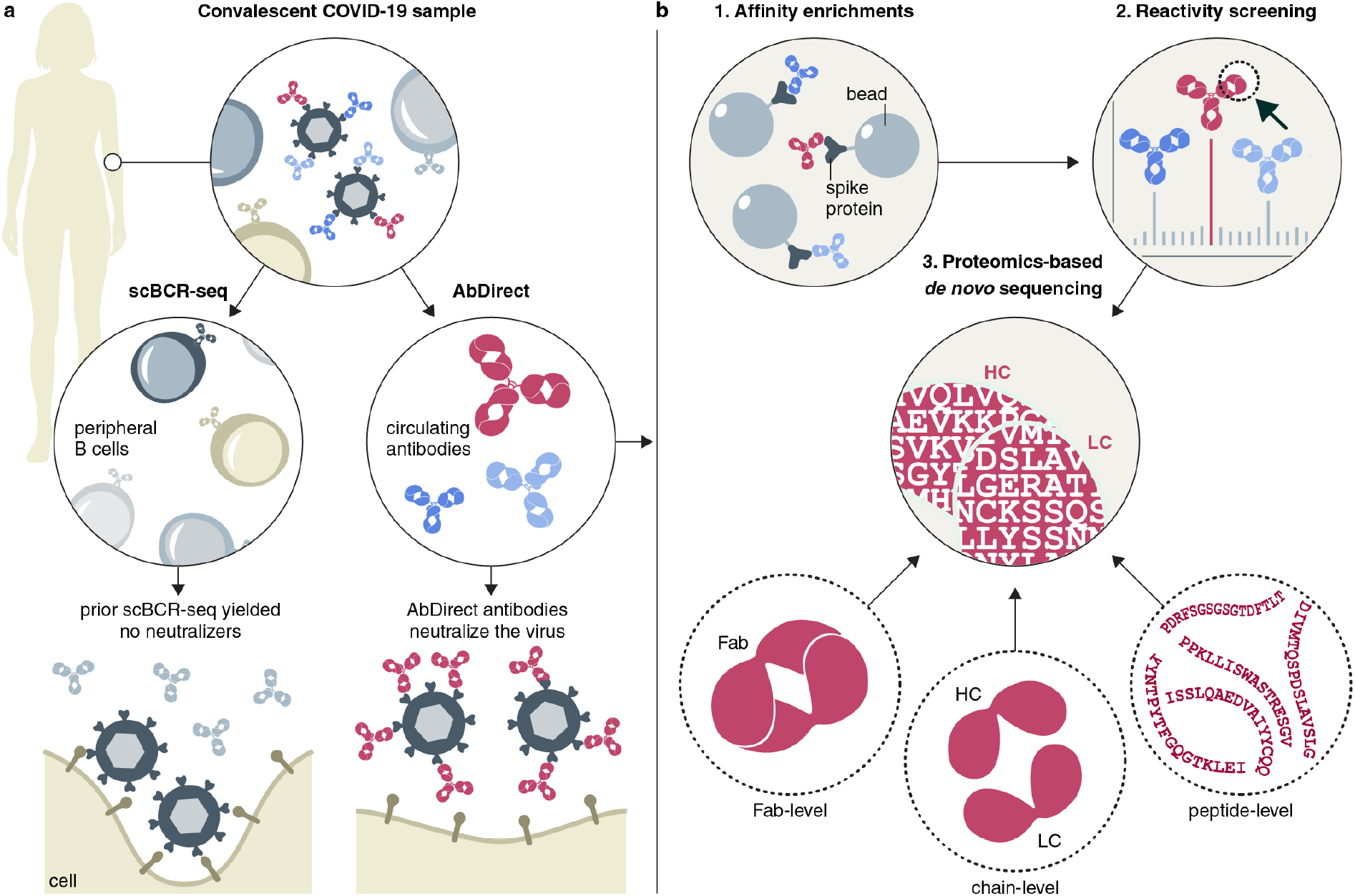
The AbDirect platform enables protein-centric antibody discovery directly from human biofluids, accessing plasma cell-secreted repertoires distinct from peripheral B cells. **a**, From the same potent convalescent COVID-19 sample where scBCR-seq previously did not identify neutralizers, we assessed whether AbDirect would enable recapitulation of the observed strong serological activity. **b**, AbDirect starts with the affinity capture of serological antibodies from plasma using SARS-CoV-2 S trimer antigens (1). Next, upfront, multidimensional screening establishes clonal-level reactivities via LC-MS antibody repertoire profiling of parallel enrichments with trimer and subunit affinity baits (2). Selected antibodies are subjected to *de novo* sequencing via integrative proteomics involving Fab-level, chain-level, and peptide-level LC-MS analyses, establishing a distinct panel of sequences (3). Resulting COVID-19 mAbs exhibited superior binding and neutralization compared to scBCR-seq outcomes, demonstrating AbDirect’s ability to access functional, *in vivo*-matured repertoires overlooked by conventional methods.

To assess serological dynamics and confirm sample selection, we first performed longitudinal antibody repertoire profiling across three sampling time points (plasma from days 23, 101, and 158 post-onset). Building further upon an approach similar to Van Rijswijck et al.^40^, we analyzed IgG1 and IgA1 Fabs obtained from whole plasma and affinity enrichments with the WT pre-fusion S protein trimer by LC-MS to detect and quantify unique Fab molecules (**Figure 2a**). In line with serum titers, day 23 showed the highest anti-S protein abundance (16% of the detected IgG1 repertoire), with dominant individual IgG1 clones reaching ∼80 µg/mL and the cumulative concentration reaching ∼172 µg/mL (**Figure 2b**). These levels declined drastically at later sampling timepoints, for some clones >10-fold (**Figure S1a**), with the fractional abundance of anti-S clones in the total detected IgG1 repertoire dropping from 16% to 9.6% to 7.4% (**Figure S1b**). Intriguingly, we observed high persistence of anti-S clones with >50% of anti-S abundance detected at day 158 representing clones already present at day 23 (**Figure 2c** and **Figure S1b**). Still, some new anti-S trimer clones appeared at later timepoints, notably several putatively Fab-glycosylated antibodies at day 158 (based on their higher mass and the monosaccharide mass differences they exhibit) (**Figure S1a**). These findings confirm that serological discovery is compatible with broad sampling windows and validate day 23 for deeper analysis, enabling a direct head-to-head comparison with the repertoire data acquired by scBCR-seq^51^.

**Figure 2.**
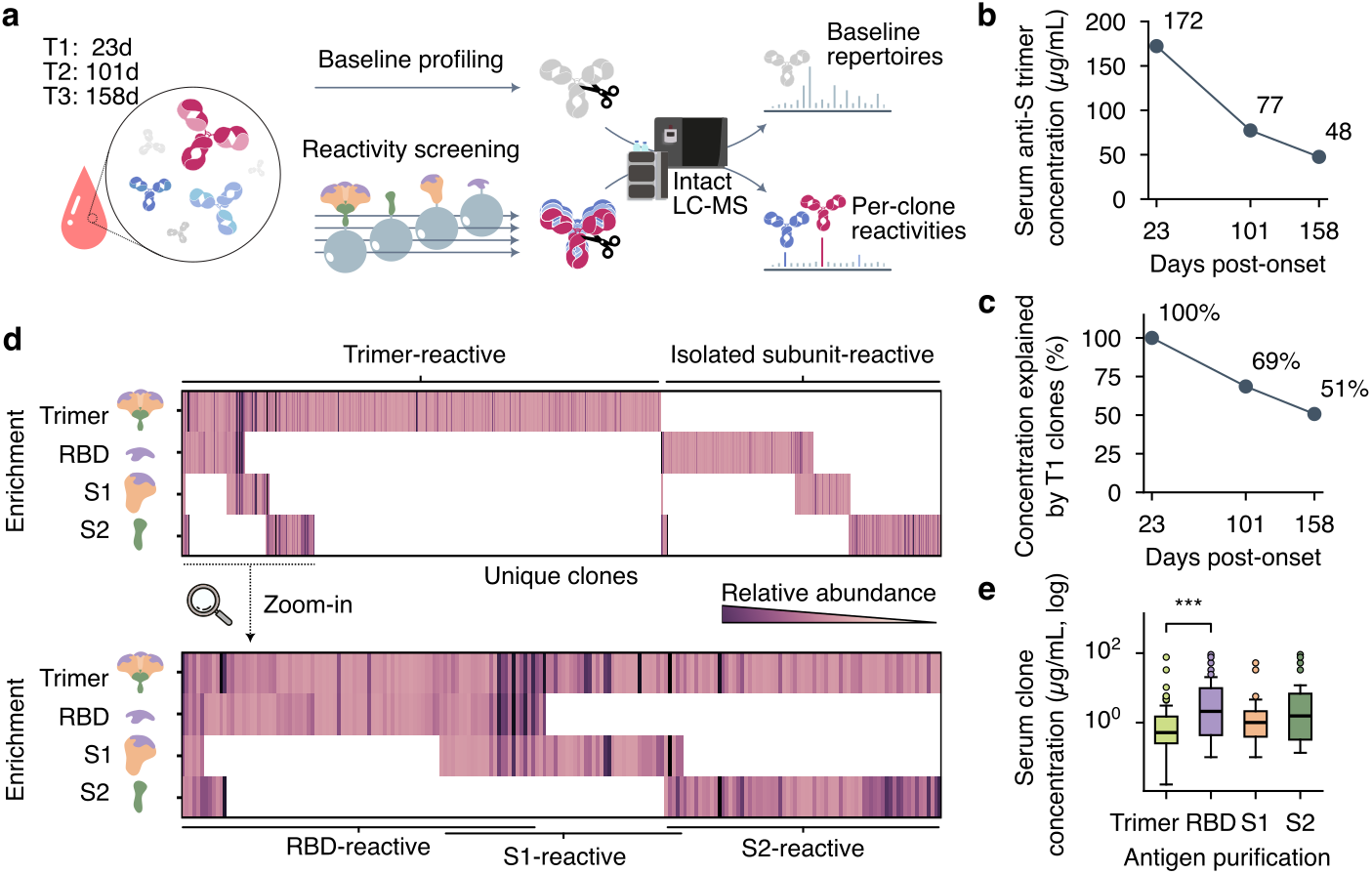
Longitudinal profiling and upfront reactivity screening reveal 1,139 distinct anti-SARS-CoV-2 S protein IgG1 clones with discrete reactivity profiles. **a**, Overview of the screening workflow, involving baseline and antigen-specific repertoire profiling of longitudinal plasma collected 23, 101, and 158 days post-symptom onset. **b**, Longitudinal cumulative plasma concentration (µg/mL) of anti-S trimer IgG1 clones across time points, highlighting a steep decline after the first collection. **c**, Percentage of the total anti-S repertoire abundance at each time point represented by clones already present on day 23 (collection 1). **d**, Heatmaps detailing reactivity profiles of the 1,139 distinct IgG1 molecules enriched from day 23 plasma with SARS-CoV-2 S protein antigens (trimers: WT and P.1, and subunits: WT RBD, S1, and S2). Clones are broadly grouped into S-trimer-reactive (n=719) and isolated subunit-only reactivities (n=420). Bottom heatmap shows a zoom-in of 200 S-trimer-reactive clones that were also detected in at least one subunit enrichment. S trimer represents the aggregate of WT and P.1 S trimer-reactive fractions. **e**, Serological abundance distribution (serum concentration in µg/mL, log scale) of IgG1 clones across different antigen reactivities, demonstrating that RBD-reactive clones show the highest average abundance (*** p < 0.001, two-sided Mann-Whitney U test comparing S trimer- and RBD-reactive clones).

### Upfront multidimensional reactivity screening identifies diverse clonal profiles

We next applied in-depth reactivity screening to the day 23 IgG1 repertoire, analyzing antibodies against a variety of SARS-CoV-2 antigens, namely the pre-fusion S trimers (WT and P.1) and several subunits (WT RBD, S1, and S2) to obtain detailed polyclonal repertoire profiles by LC-MS (**Figure 2a**). This high-throughput, multidimensional approach required only 20 µL of plasma for each enrichment and detected 1,139 distinct anti-S protein IgG1 molecules, spanning a wide dynamic range (>4 orders of magnitude in abundance), highlighting the enormous antigen-specific serological repertoire diversity already in a single donor (**Figure 2d**). Notably, the serological abundance of isolated RBD-reactive clones was highest of all reactivities, even higher than intact S trimer-reactive clones (**Figure 2e** and **Figure S2**). This corroborates the immunodominant nature of the RBD and demonstrates AbDirect’s ability to directly quantify clonal dominance in both serum and in enriched fractions – information that is not easily captured by scBCR-seq.

Within this dataset, several distinct antibody reactivity clusters could be discerned. Expectedly, repertoires targeting S1 and S2 showed minimal overlap, whereas 76% of RBD-reactive abundance cross-reacted with S1 (**Figure 2d** and **Figure S2b**). Interestingly, roughly 48% of RBD- and S2-reactive abundance was not detected in the full S trimer-based enrichments, suggesting cryptic or post-fusion epitopes. Conversely, about 28% of trimer-reactive abundance, was not reactive with the free individual subunits, indicating trimer-specific or quaternary epitopes. These seemingly disparate epitope patterns underscore the value of upfront screening to prioritize clones. Cross-reactivity analysis furthermore revealed that about 53% of anti-WT S trimer-reactive clones also bound the P.1 variant trimer, explaining almost all abundance in the P.1-based enrichment. However, some of these cross-reactive clones were detected at lower abundance, suggesting weaker interactions (**Figure S2**). Clones within this cross-reactive fraction targeted both S1 and S2 subunits, with the notable exception of the RBD, likely reflecting the extensive mutations in this domain in the P.1 variant from WT. Overall, this upfront clonal-level reactivity profiling thus enables the informed inclusion and exclusion of samples and clones for further analysis, potentially reducing downstream validation and selection efforts.

### Integrative proteomics enables high-resolution *de novo* sequencing

Sequential affinity enrichments (using P.1 (gamma) and subsequently WT trimer as antigen) were next subjected to an integrative proteomics workflow for sequence discovery to study cross-reactive and WT-specific clones in more detail (**Figure 3a, b** and **Figure S3**). Building on prior methods reported by Bondt et al.^32^ and Schulte et al.^52^, this approach combined *de novo* peptide sequencing by using a multi-protease bottom-up proteomics approach with protein-centric electron transfer dissociation (ETD) MS2 of intact reduced chains and Fab molecules. To resolve chain pairings in complex polyclonal samples, candidate matches between chains and parent Fabs were evaluated against orthogonal LC-MS attributes and manually validated to identify the best match. Resulting per-clone, multi-layer data bundles were generated as input for targeted *de novo* sequencing. In brief, putative variable gene segments (consensus) were first assembled from bottom-up *de novo* peptide-spectrum matches (PSMs) using human germline sequences as templates. Fab- and chain-level ETD MS2 data bundles (scaffolds) were matched to these putative consensus sequences. Possible mutations were then iteratively processed via gap filling of the scaffold and ambiguity resolution of the consensus until full agreement was reached between all three layers of MS evidence (see method overview in **Figure 3a** and an example of clone-specific sequencing dataset in **Figure S4**). Combined, this enabled the refinement of true and complete individual serological antibody sequences rather than consensus sequences, directly from complex polyclonal samples and using minimal amounts of materials (1-5 µg per affinity-enriched IgG1/IgA1 fraction).

**Figure 3.**
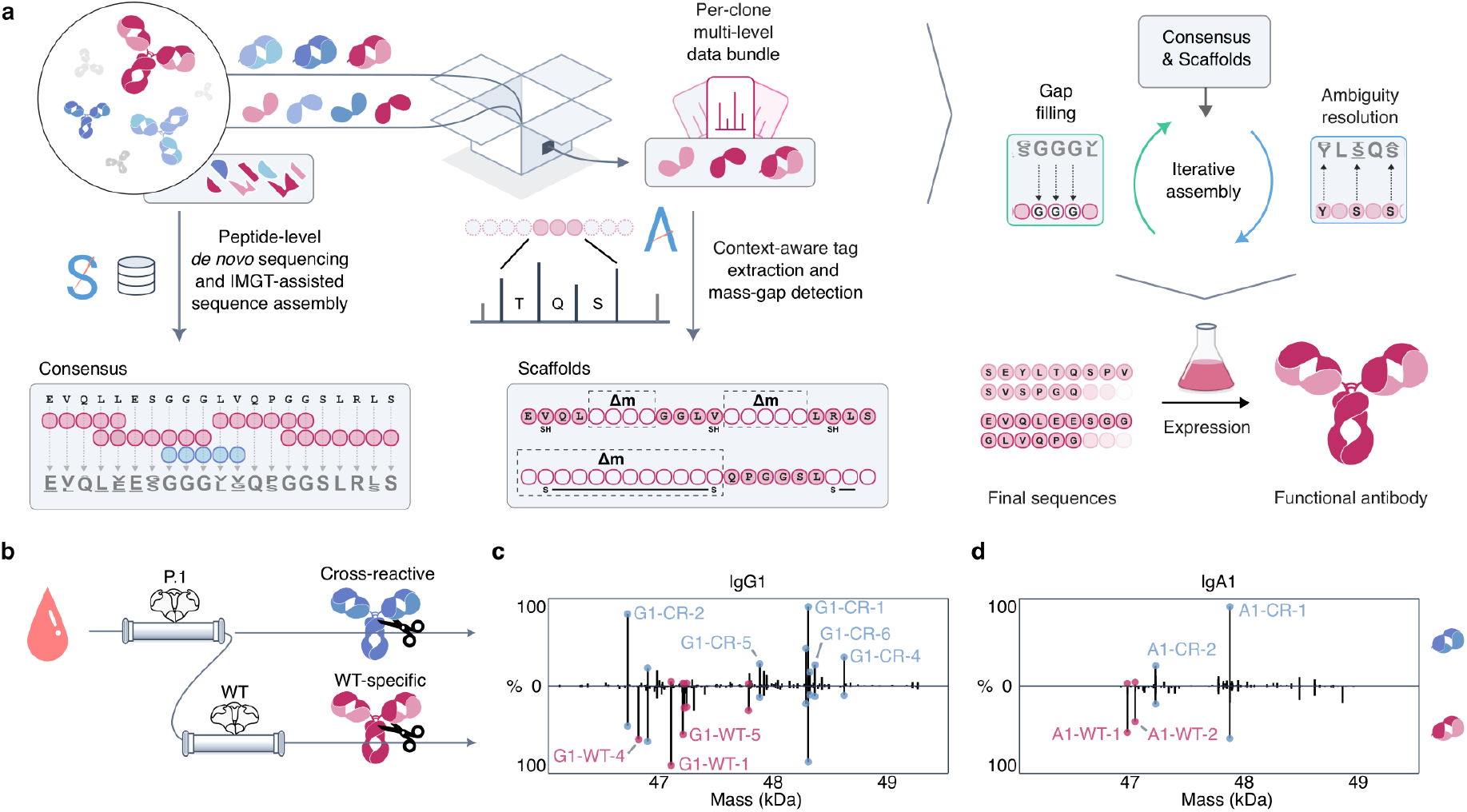
Integrative proteomics enables the discovery of 14 IgG1 and 4 IgA1 chain-paired antibody sequences against the SARS-CoV-2 S protein from COSCA3 T1 plasma. **a**, Overview of integrative mass spectrometry-based *de novo* sequencing workflow for polyclonal antibody samples. Three MS data streams (peptide-, chain-, and Fab-level) are processed to provide consensus sequences and scaffolds, which are then iteratively assembled into full, chain-paired sequences for functional antibody expression. **b**, Overview of the sequential affinity enrichment strategy to isolate cross-reactive (P.1) and WT-specific S protein-reactive clones. **c**, Mirrored IgG1 Fab mass plots for the cross-reactive fraction (top) and the WT-specific fraction (bottom). Sequenced clones are labelled with colored dots corresponding to the fractions in **b. d**, Mirrored IgA1 Fab mass plots for cross-reactive and WT-specific enrichments. Several representative anti-S protein antibody clones are annotated in the mirror plots. See **Figure S5** for the full list of discovered clones.

This standalone approach – requiring no parallel BCR-seq data – resolved a panel of full-length, chain-paired sequences directly from complex polyclonal mixtures. It enabled us to successfully sequence all clones at >10% relative abundance in the affinity-enriched IgG1 and IgA1 fractions, yielding 18 antibody sequences (**Figure 3c, d**, and **Figure S5**). Notably, these also included four IgA1 clones, a class rarely captured by conventional scBCR-seq pipelines that focus primarily on IgG or IgM. The platform proved able to resolve complex polyclonal mixtures of hundreds of antibodies, identifying individual sequences and distinguishing minute differences between clones. For example, we observed that several clones shared an identical heavy or light chain but paired differently. Near-identical variants could also be discerned, such as clones G1-CR-6 and G1-CR-8, differing by one amino acid in the light or heavy chain from G1-CR-1. Another clone appeared both as IgA1 and subsequent class-switched IgG1 without further maturation. Intriguingly, clone G1-CR-2 was found to harbor two extra cysteines (A24C mutation in FR1 of the IGHV3-48 HC and germline C32 in FR2 of the IGLV3-1 LC), inducing the formation of alternative disulfide bonding with an additional interchain disulfide, which was corroborated by distinctive fragmentation patterns in the ETD MS2 analysis of the Fab. Together, these features illustrate the resolving power of the approach in sequence discovery from real-world endogenous samples.

### Serological antibodies represent a distinct population not represented by captured peripheral memory B cells

Comparison of the serological panel to the scBCR-seq dataset of 105 unique VH and 27 unique VL sequences from exactly the same sample revealed a profound divergence. Not one direct or near-identical paired match could be found, with even the closest match (clone A1-CR-2 with BCR-derived COVA3-10) sharing only 80/92% VH/VL identity. Single-chain matches were similarly non-existent, with the closest matches differing by at least one amino acid. While deeper sampling might have uncovered more overlap, these findings corroborate that dominant serological antibodies represent a distinct population that is not necessarily captured by genetic sequencing of peripheral memory B cells. This highlights the unique value of direct protein-level discovery to identify the most dominant circulating clones.

Clonal lineage analysis placed serological and scBCR-seq sequences mostly in separate clusters, though spanning similar overall diversity (**Figure 4a** and **Figure S6**). When considering confirmed binding scBCR-seq branches only, sequences clustered closer together, but large low-binding scBCR-seq clusters did not have serological counterparts. IGHV4-39 stood out as a prevalent germline in both datasets, representing 22% and 25% of protein and BCR-derived sequences. Beyond that, germline usage differed markedly between the datasets (**Figure 4b**). For example, IGHV1-46 and IGLV10-54 (rare in CoV-AbDab^53^, ∼0.05%) were enriched in the serological panel but absent in scBCR-seq. Conversely, scBCR-seq overrepresented IGHV3-7, IGHV3-NL1, IGKV1-16, and IGKV3-20, which were not among the sequenced, highly abundant affinity-enriched clones in the here presented serological data (**Figure S6**). Furthermore, while heavy chain CDR3s were of similar length in both datasets, serological light chain CDR3s were longer (∼1 amino acid, p=0.0012, **Figure 4c**). Overall somatic hypermutation rates from the germline were comparable (**Figure 4d**), but serological clones showed higher maturation *versus* a subset of 10 confirmed scBCR-seq binders. Importantly, serological sequences were more similar to confirmed binders and neutralizers in the public CoV-AbDab, especially in their CDR3s (**Figure 4e**). Taken together, these data indicate that direct serological discovery targets a distinct immunological population and may deliver more relevant antibody sequences.

**Figure 4.**
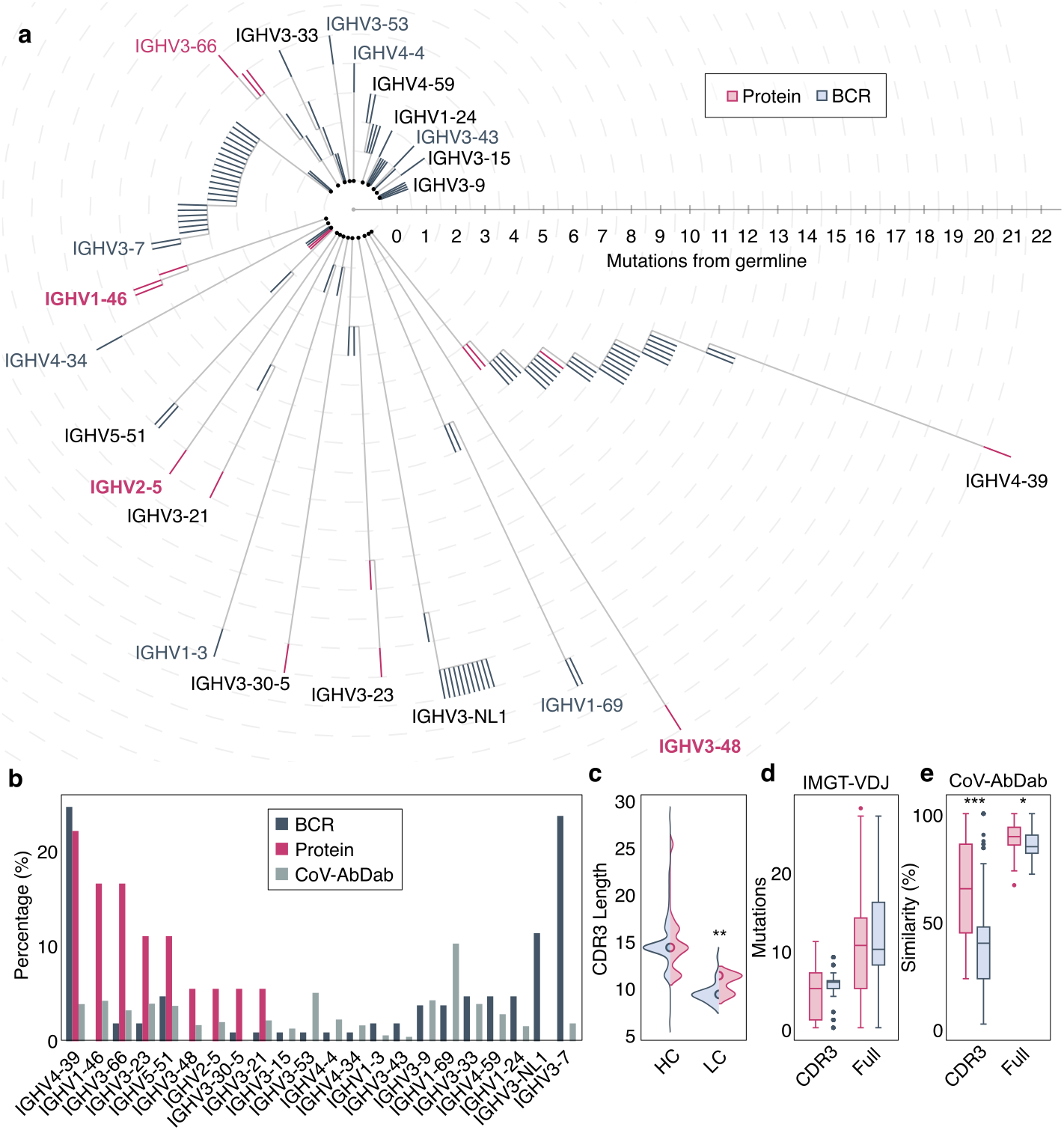
Distinctive genotypic signatures of serological antibody and BCR sequences against the SARS-CoV-2 S protein. **a**, Germline-centered clonal lineage tree of 18 protein- and 105 BCR-derived VH regions showing somatic hypermutation-based clonal relationships. Each concentric ring represents one amino acid substitution from the germline V-gene (center). Crimson-colored leaves represent serological sequences, steel-colored leaves represent BCR sequences. VH region germlines are indicated near the most mutated clone, with protein-only groups in bold crimson and unproductive BCR-only groups in steel color. **b**, Distribution of VH germline usage of serological antibody (crimson), BCR (steel), and human CoV-AbDab^53^ (gray) sequences. Only germlines present in at least two groups are shown. **c**, Distribution of CDRH3 and CDRL3 lengths of serological antibody (crimson) and BCR (steel) sequences. Half-dot represents the median value for each group. **d**, Boxplots of absolute mutation rates in the sequence segments of serological antibody (crimson) and BCR (steel) sequences compared to the closest matching germline, showing similar overall mutation rates. **e**, Boxplots of similarities between the serological antibodies (crimson), BCR (steel) sequences and the closest matching binding/neutralizing VH/VL sequences from the CoV-AbDab, with protein-derived sequences showing higher similarity, especially in CDR3. Pairwise statistical differences were assessed using the two-tailed Mann-Whitney U test and annotated where significant (significance levels: * p < 0.05; ** p < 0.01; *** p < 0.001).

### Recombinant serological antibodies demonstrate superior binding and neutralization

All 18 discovered serological antibody sequences were successfully expressed in IgG1 format (100% expression rate), all yielding soluble protein suitable for downstream characterization. Luminex binding assays confirmed that 15 out of 18 antibodies (83%) strongly interacted with the S protein trimer, exhibiting nM to pM range EC_50_ values (**Figure 5a** and **Figure S7**). Most clones enriched with the P.1 variant cross-reacted with more recent variants, including JN1 (Omicron lineage), and in agreement with upfront reactivity screening data, these did not bind the RBD. Conversely, most WT-enriched clones were confirmed to target the RBD and did not cross-react beyond the Delta variant. Notably, the serological approach yielded multiple high-affinity clones with sub-nanomolar apparent affinities, contrasting sharply with the prior scBCR-seq panel, which lacked comparable high-affinity interactors.

**Figure 5.**
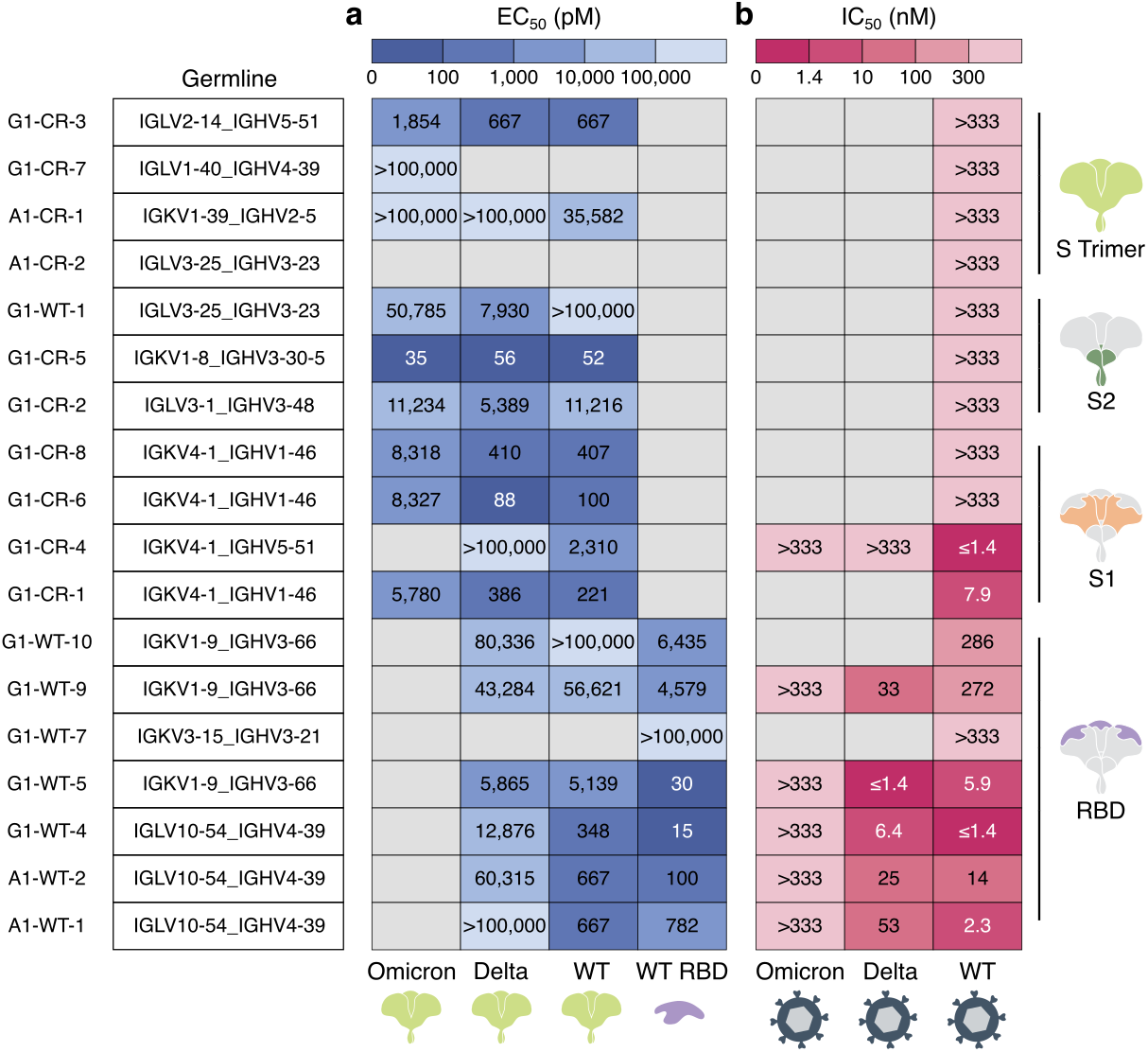
Functional validation of serological SARS-CoV-2 S protein-specific mAbs discovered from COSCA3 T1 plasma. **a**, Heatmap showing binding EC_50_ values in pM as determined from Luminex bead-based assays. **b**, Heatmap showing neutralization IC_50_ values in nM as determined from pseudovirus neutralization assays. Gray-shaded cells indicate cases where IC_50_ or EC_50_ values could not be reliably determined. Cartoons on the right indicate most likely antigenic clustering as determined by upfront reactivity screening (see **Figure 2**). The germline represents the closest matching VH and VL alleles for each clone.

To functionally validate the AbDirect-derived mAbs, we performed well-established pseudovirus neutralization assays using WT, Delta, and XBB.1 (Omicron lineage) variants^54,55^. This identified 8 out of 15 binding antibodies (53%) as genuine neutralizers, including predominantly clones targeting RBD, but also two S1-specific antibodies (**Figure 5b**). Five clones demonstrated high potency (IC_50_ < 10 nM), of which three were of very high potency (≤ 1.4 nM, lowest concentration tested). Intriguingly, S1-targeting clones that cross-bound JN1 neutralized only the WT variant, whereas some less cross-reactive clones targeting RBD also neutralized the Delta variant. These neutralization results contrast with the prior scBCR-seq panel from this donor, in which the ten clones tested did not include any neutralizers. AbDirect thus recapitulated the observed serum neutralization activity, yielding multiple potent antibodies from a single donor for which prior scBCR-seq had been unsuccessful. In summary, of the 18 antibodies sequenced, all 18 (100%) expressed, 15 (83%) bound the S protein, 8 (44%) neutralized SARS-CoV-2 pseudovirus, 5 (28%) did so potently (IC_50_ < 10 nM), and three highly potently (IC_50_ ≤ 1.4 nM). AbDirect thereby establishes a clean conversion funnel from polyclonal serum to functionally validated leads.

## Discussion

In the proof-of-concept study described here, we provide evidence that proteomics-based serological antibody discovery and scBCR-seq sample immunologically distinct populations with divergent sequences and functional profiles. This was highlighted by the absence of any near-identical paired matches, marked differences in germline representation, and closer alignment of protein-derived sequences to public, validated COVID-19 antibodies in the CoV-AbDab^53^. Notably, AbDirect enabled us to discover multiple high-potency IgG1s and, importantly, IgA1s from a convalescent sample with strong serum neutralization, outperforming prior results obtained via established scBCR-seq approaches^51^. Extending beyond prior proteogenomic studies^16,17,34,44,47^, these findings underscore that antibodies secreted by plasma cells are not reliably captured from peripheral memory B cells, highlighting the value of protein-centric discovery for accessing *in vivo*-matured repertoires that drive humoral protection.

The technical strengths of the AbDirect platform further empowered these insights. Upfront multidimensional reactivity screening enabled the discernment of diverse clonal profiles, including trimer-specific, cryptic epitope-targeting, and cross-reactive subsets. Furthermore, it uniquely captures the serological abundance of each clone and identifies the most dominant functional clones – a critical feature not offered by scBCR-seq. This allows rational prioritization of samples and antibodies before mass spectrometry-based *de novo* sequencing, potentially reducing downstream validation efforts compared to conventional genetic approaches. Through integrative proteomics, we achieved high-resolution standalone *de novo* sequencing, even in complex polyclonal mixtures with hundreds of unique molecules. This allowed us to resolve variants differing by single amino acids, class-switched chain pairs, and unusual structural features such as additional disulfides in the variable region. The platform enabled standalone operation with minimal polyclonal input (e.g., 1-5 µg affinity-enriched IgG1/IgA1 from 1.5 mL of plasma), while preserving qualitative and quantitative information on the native polyclonal context and isotype diversity.

Taken together, these advancements position serological proteomics as a compelling tool to complement scBCR-seq, which so far has remained dominant due to its higher throughput, lower cost, and proven success in identifying potent antibodies across many applications^5,18–21^. While scBCR-seq excels in samples with relevant viable cells, protein-centric serological discovery specifically addresses important gaps. These include accessing mature antibodies actively secreted in circulation, also from archived biofluids, which is important in cases where peripheral B cells are inaccessible or do not represent the functional response^18,22,23,39^. Extending beyond foundational integrated proteomics^32,46^ and hybrid proteogenomics^5,34,35,44^, AbDirect demonstrates enhanced yields, sensitivity, and resolution without reliance on parallel DNA/RNA sequencing. Furthermore, in the future it may also be integrated with cryo-EMPEM of the same polyclonal samples, enabling simultaneous high-resolution sequence discovery and structural epitope mapping on native antibodies^56^.

As a first benchmark on a single COVID-19 donor, there are still limitations to this study, and generalizations warrant some caution. Firstly, both discovery approaches were limited in sampling depth, and more overlap between serological and peripheral BCR repertoires may emerge with deeper sampling. We also cannot exclude that broader expression of the scBCR-seq panel would have identified neutralizers, though the original prioritization by S-protein affinity represents the standard approach. Findings furthermore reflect a severely ill, early convalescent donor with abundant anti-S protein clones, and broader validation across diverse individuals and diseases is needed. Current proteomics methods furthermore favor dominant clones in high-titer samples, with sensitivity gains or larger samples required for reliable sequencing of lower-abundance antibodies. As a novel approach, laboriousness and cost also remain high compared to scBCR-seq.

Importantly, this work validates serological discovery for high-value scenarios, including retrospective biobank studies and cases where more invasive scBCR-seq is not feasible or underperforms. Beyond infectious diseases, the platform holds promise for autoimmune and neurodegenerative disorders, where direct access to autoantibodies could benefit the field^19,57^. In oncology, tumor-specific human antibodies recovered directly from patients could provide powerful starting points for antibody-drug conjugates, bispecific T-cell engagers, or CAR-T constructs. Future improvements in sensitivity and throughput will broaden these applications, complementing genetic methods for mapping humoral immune responses and accelerating therapeutic lead discovery. In conclusion, this study thus establishes AbDirect as a powerful tool for delivering *in vivo*-matured antibody sequences directly from serum, complementing existing cell-based discovery approaches by unlocking previously overlooked antibodies in functional repertoires.

## Materials & Methods

### Donor plasma and antibody enrichment

Convalescent plasma from donor COSCA3 (early pandemic COVID-19; collection 23, 101, and 158 days post symptom onset) was obtained from the COSCA and RECoVERED studies (see Ethics Statement)^51^. SARS-CoV-2 S protein-reactive antibodies were affinity captured from plasma by magnetic beads coupled to prefusion-stabilized S trimers (WT or P.1) or isolated subdomains (WT RBD, S1, S2; all ACRObiosystems), using protocols adapted from Van Rijswijck et al.^40^. For *de novo* sequencing, antibodies were enriched sequentially from ∼2 mL plasma using P.1 S trimer beads, followed by loading of the unbound fraction onto WT S trimer beads (yielding ∼5-10 µg of enriched immunoglobulin per variant).

### Fab fragment generation

IgG1 and IgA1 Fabs were generated from plasma and enriched fractions as described previously^40,58^. IgG and IgA fractions were purified sequentially using CaptureSelect Fc-XL and IgA-XL affinity matrix (Thermo Scientific). IgG1 Fabs were generated with FabDELLO, IgA1 Fabs with OpeRATOR and SialEXO (both Genovis). Fc and undigested antibody fractions were removed by re-affinity capture, and flow-throughs containing Fabs were collected. Internal standard mAbs were spiked before digestion for normalizing abundance (IgG1: trastuzumab, alemtuzumab; IgA1: 7D8, 5D5).

### Multi-protease bottom-up sample preparation

Purified Fab fractions were processed for multi-protease digestion by the SP3 protocol^59^. After reduction with TCEP and alkylation with iodoacetic acid, each sample was split into five portions and digested overnight at 37 °C using trypsin, chymotrypsin, elastase, alpha-lytic protease, or Lys-C (1:25 m/m).

### LC-MS for Fab-level, reduced chain-level, and peptide-level analyses

Fabs and reduced chains (TCEP) were analyzed by reversed-phase LC-MS on a Vanquish Neo UHPLC coupled to an Orbitrap Eclipse Tribrid mass spectrometer (Thermo Fisher Scientific), using methods adapted from Bondt et al.^32^. Separation was performed on a MAbPac Capillary RP column (150 µm × 150 mm) at 80 °C. For repertoire profiling, MS1 data were acquired only. For sequencing, ETD MS2 spectra were additionally acquired for intact Fabs and reduced chains. Digested peptide mixtures were separated on an UltiMate 3000 UHPLC (Thermo Scientific) using a custom C18 column (Poroshell 120 EC-C18, 2.7 µm, 50 cm × 75 µm, Agilent) using methods adapted from Peng et al.^59^. Precursors were fragmented using stepped HCD (NCE 22/28/34%) or EThcD (27% supplemental NCE), with MS2 acquired at 30,000 resolution.

### Antibody repertoire profiling

Fab-level MS1 data were deconvoluted in BioPharma Finder v5.1 (ReSpect algorithm; Thermo Fisher Scientific) and analyzed in a Python 3.13 notebook adapted from Bondt et al.^32^. Briefly, antibody clones were defined by unique mass and retention time combinations, quantified by normalization to internal standard mAbs, and matched between samples by UPGMA hierarchical clustering using L∞ distance. For antigen reactivity profiling, complete linkage clustering with correlation distance was applied to log_10_-transformed relative abundances.

### Integrative *de novo* antibody sequence assembly

Chain-paired, full variable region antibody sequences were assembled by integrative proteomics, building on Bondt et al.^32^ and Schulte et al.^52^. Bottom-up data were processed for *de novo* peptide sequencing in PEAKS Studio v11.5 (precursor 20 ppm, fragment 0.02 Da; fixed: carboxymethylation of Cys; variable: carboxymethylation of N-termini/Lys, oxidation of Met/Trp, pyroglutamic acid from N-terminal Glu/Gln). Peptides were assembled into gene segment libraries using Stitch v1.5.0^52^ with the IMGT human germline database^60^ as templates. Per-clone datasets were assembled by integrative matching of Fab-level, chain-level, and peptide-level MS data based on orthogonal LC-MS attributes and manual validation of putative chain pairings. Briefly, V-gene assemblies from Stitch were iteratively refined against ETD MS2 data from reduced chain pairs using Annotator^61^, and CDR3/J segments were further resolved using intact Fab ETD MS2. Leucine/isoleucine assignments were made based on *w*-ions in EThcD peptide data^62^. Final sequences were confirmed by manual inspection across all data layers.

### Genetic analysis of serological and BCR repertoires

BCR sequences from COSCA3 (105 VH and 27 VL) were obtained from the Brouwer et al. study^51^. All sequences were annotated with IgBLAST v1.22.0 against IMGT V(D)J germlines^60^, CoV-AbDab^53^ (COVA3 sequences excluded), and the COSCA3 BCR sequences. Mutation rates were assessed using Align-CLI, allowing up to two isobaric substitutions to account for any MS-derived ambiguities. CDR3 length and mutation rate differences between serological antibodies and BCRs were evaluated by two-sided Mann–Whitney U test (SciPy). Clonal expansion across germline V genes (Figure 4a, Figure S6c) was visualized as a germline-anchored radial diagram, with each sequence positioned radially according to SHM count. A complementary UPGMA similarity dendrogram across all serological, BCR, and germline-match sequences (Figure S6a, b) was built from pairwise amino-acid Hamming distances using average-linkage clustering (SciPy) with SHM-proportional leaf extensions.

### Recombinant monoclonal antibody expression

Variable region sequences were cloned into IgG1 expression vectors and expressed in HEK293F cells as previously described^63,64^. Antibodies were purified by Protein G affinity chromatography, concentrated, and buffer-exchanged to PBS using 100 kDa MWCO spin filters. Protein concentrations were determined by UV280.

### SARS-CoV-2 spike protein expression

Spike and RBD variants were transiently expressed in HEK293F cells and purified by Strep-TactinXT affinity chromatography as previously described^51,65^.

### Luminex binding assay

Antigen-specific binding was assessed by Luminex, adapted from Grobben et al.^66^. S protein antigens were covalently coupled to MagPlex beads by two-step carbodiimide chemistry. Antibodies were tested across a concentration range of 7 fM to 70 nM. EC_50_ values were determined by Michaelis– Menten fitting or, where R^2^ < 0.9, four-parameter logistic regression.

### Pseudovirus neutralization assay

Neutralization was assessed using a SARS-CoV-2 pseudovirus assay as previously described^54,55^. Pseudoviruses were produced in HEK293T cells using pHIV-1NL43 ΔEnv-NanoLuc, and neutralization was measured against HEK293T/ACE2 cells by NanoLuc luciferase readout. IC_50_ values were determined by nonlinear regression in GraphPad Prism.

## Ethics Statement

This study used convalescent plasma from donor COSCA3, originally collected as part of the COSCA (protocol NL73281.018.20) and RECoVERED (protocol NL73759.018.20) cohort studies at Amsterdam University Medical Centers, location AMC, the Netherlands^51^. Both studies were approved by the local medical ethics committee of the Amsterdam UMC. Written informed consent was obtained from all participants prior to enrolment. No new participants were recruited, and no new samples were collected for the current study.

## Data Availability

Mass spectrometry raw data, peptide identification results, and antibody sequences supporting the findings of this study will be deposited in public repositories at the time of journal publication. The mass spectrometry data will be deposited in the ProteomeXchange Consortium via the PRIDE partner repository (https://www.ebi.ac.uk/pride/). All other data supporting the findings of this study are available from the corresponding author upon reasonable request.

## Code Availability

The results in this study were generated using openly available, cited software (PEAKS Studio v11.5, Stitch v1.5.0, Annotator). Code for downstream data analysis and figure generation will be made available with the peer-reviewed publication of this work.

## Acknowledgements

This work was principally supported by NWO Take-off Phase 2 innovation loan #22014 to MAdB/Abvion. AJRH acknowledges funding from the ERC Advanced Grant #101141457 and NWO Spinoza Prize SPI.2017.028, and additional support from the Large-Scale Research Infrastructure - National Roadmap grant BioBeyond_NL (184.037.015) and PharmaNL grant “PharmIgA”. The authors thank all members of the Biomolecular Mass Spectrometry & Proteomics group at Utrecht University, in particular Arjan Barendregt and Peter Mosen for experimental support and Albert Bondt and Joost Snijder for fruitful scientific discussions. The authors thank all members of the Department of Medical Microbiology and Infection Prevention at Amsterdam UMC, particularly Rogier W. Sanders and Tom G. Caniels, for supporting the antibody production and validation, and Maria Prins and Menno D. de Jong for providing longitudinal samples from the RECoVERED cohort. The authors thank patient COSCA3 for participating in the COSCA and RECOvERED studies. The authors thank Genmab for providing internal reference mAbs to AJRH and for supporting Abvion’s NWO Take-off Phase 2 application. The authors thank Joost Bakker at Scicomvisuals for visual design support, particularly for Figure 1.

## Author contributions

Conceptualization and methodology: MAdB, ST, AJRH, and MJvG. Software: ST and DS. Investigation and validation: ST, MAdB, JvR, JAB, DMHvR, and MM. Formal analysis and data curation: ST, MAdB, DMHvR, and JvR, supported by DS. Resources: MAdB, AJRH, MJvG, and GJdB. Writing-original draft: MAdB and ST. Writing-review & editing: MAdB, ST, AJRH, MJvG, and DMHvR with input from all listed authors. Visualization: ST and MAdB. Supervision: MAdB, AJRH, and MJvG. Funding and administration: MAdB, AJRH, and MJvG.

## Competing interests

ST, MAdB, and AJRH are employees or contractors of Abvion, hold equity interests in Abvion, and are inventors on a patent application related to the antibody discovery platform. MM is an employee of Abvion. Abvion has a material transfer agreement with Amsterdam UMC, employer of MJvG, JvR, JAB, and GJdB. Abvion is commercializing the antibody discovery platform described in this study. Remaining authors declare no competing interests.

## Footnotes

a Plasma was used as the biofluid throughout; “serum” is used per conventional serological terminology.

