## Supplementary Information for "Direct serological antibody discovery by integrative proteomics yields potent neutralizers overlooked by single-cell BCR sequencing"

### Supplementary Materials and Methods

#### Affinity Capture and Preparation of Antibodies

Convalescent plasma from donor COSCA3 (early pandemic COVID-19; collection from 23, 101, and 158 days post symptom onset) was obtained from the COSCA and RECoVERED studies (see Ethics Statement)[1]. Antibodies against SARS-CoV-2 spike (S) proteins were enriched through affinity capture using magnetic beads with prefusion-stabilized SARS-CoV-2 S trimer variants (WT or P.1, chosen for commercial availability; ACRObiosystems) or isolated domains (WT RBD, S1, S2; ACRObiosystems), following protocols adapted from Van Rijswijck et al.[2]. Beads were washed and blocked with PBS containing 0.05% (w/v) BSA and 0.05% (v/v) Tween-20 (PBS-BSA-Tw) before use. Plasma was diluted 10-fold in the same buffer before loading. Incubations were performed overnight at 7 °C on a tube rotator. Following capture, the unbound fraction was stored, and beads were washed three times with PBS-BSA-Tw and twice with MilliQ water + 0.05% Tween-20. Bound antibodies were eluted with 0.4% formic acid + 0.05% Tween-20, followed by immediate neutralization with 1 M Tris pH 8.5 at a 1:4 (v/v) ratio. Variations in scale, antigen selection, and enrichment strategy were applied depending on the experimental purpose, as detailed below.

For repertoire profiling, enrichments were performed in parallel using 100 µg of magnetic beads per antigen. For initial longitudinal profiling, approximately 5 µL of plasma (diluted 10-fold) was incubated with WT SARS-CoV-2 S trimer beads only. For reactivity profiling, enrichments were similarly conducted in parallel using 20 µL of plasma (diluted 10-fold) incubated separately with WT or P.1 S trimer, or WT RBD, S1, or S2 domain beads. For larger-scale enrichments intended for sequencing, antibodies were captured sequentially to maximize yield. Approximately 2 mL of plasma (diluted 10-fold) was first loaded onto 2 mg of P.1 S trimer beads. Unbound antibodies from this step were then transferred to 2 mg of WT S trimer beads. Yields were approximately 5-10 µg of enriched immunoglobulin material per variant.

#### Generation of IgG1 and IgA1 Fab fragments

Plasma, depleted plasma, and enriched antibody fractions were processed to generate IgG1 and IgA1 Fab fragments for repertoire profiling and sequencing, following protocols adapted from earlier reported methods[2]. For longitudinal profiling, internal standard mAbs (IgG1: Trastuzumab, Alemtuzumab; IgA1: 7D8, 5D5) were spiked into plasma at 20 µg/mL or in enriched fractions at 2 µg/mL and used to normalize quantification.

CaptureSelect Fc-XL (IgG) or IgA-XL (IgA) affinity matrices (Thermo Scientific) in spin columns were pre-washed with PBS-BSA-Tw, after which plasma, depleted plasma, and enriched antibody fractions were loaded. IgG purifications were incubated for 60 min at room temperature on a thermomixer (800 rpm), with unbound flow-through subsequently loaded onto IgA-XL columns for similar incubation. After incubation, columns were washed three times with PBS and twice with MilliQ water. Bound antibodies were eluted three times with 0.1% formic acid and neutralized with 300 mM Tris + 900 mM NaCl + 60 mM CaCl2 pH 7.6 (1:5 v/v).

IgG1 Fabs were generated by digesting the IgG fraction with FabDELLO (Genovis) overnight at 37 °C on a thermomixer (800 rpm). IgA1 Fabs were similarly generated by digesting the IgA fraction with OpeRATOR and SialEXO (Genovis). Digested samples were reloaded onto their respective affinity matrices (neutralized with MilliQ water) and incubated for 60 min at room temperature (800 rpm) to capture Fc portions and intact antibodies. Fab molecules in the flow-through were collected for downstream profiling or sequencing.

#### Multiple Protease Digestion for Bottom-Up Proteomics

Purified IgG1 and IgA1 Fab molecules against WT and P.1 SARS-CoV-2 S trimers were processed separately for multi-protease digestion using single-pot solid-phase-enhanced sample preparation (SP3), adapted from Peng et al. (2021)[3]. Briefly, antibody samples were denatured, reduced with 10 mM tris(2-carboxyethyl)phosphine (TCEP), and alkylated with 40 mM iodoacetic acid. Each sample was divided into five equal portions and loaded onto a 1:1 mixture of hydrophilic and hydrophobic Sera-Mag SpeedBead carboxylate-modified magnetic beads (Cytiva). Beads were washed to remove contaminants, and each portion was digested overnight at 37 °C on a thermomixer (1000 rpm) with one of the following proteases at a 1:25 enzyme-to-protein ratio (m/m): trypsin, chymotrypsin, elastase, alpha-lytic protease, or Lys-C. Digestion was quenched by acidification with 1% trifluoroacetic acid (TFA), and peptides were captured.

#### LC-MS Analysis of Digested Peptides

Peptide mixtures from each digestion were analyzed separately by reversed-phase liquid chromatography-mass spectrometry (LC-MS) using an UltiMate 3000 UHPLC system coupled to an Orbitrap Eclipse Tribrid mass spectrometer (both Thermo Scientific). Chromatographic separation was performed on a custom C18 column (Poroshell 120 EC-C18, 2.7 µm particles; 50 cm × 75 µm i.d.; Agilent Technologies) at 40 °C with a flow rate of 0.3 µL/min. Mobile phase A was acidified MilliQ water, and mobile phase B was acidified acetonitrile. MS1 spectra were acquired over the m/z range of 350-2000 at a resolution of 60,000, with an automatic gain control (AGC) target of 100% and a maximum injection time of 50 ms. Precursors were isolated using a 1.6 m/z window and fragmented using either stepped high-energy collision dissociation (sHCD) at normalized collision energies (NCE) of 22%, 28%, and 34%, or electron-transfer/higher-energy collision dissociation (EThcD) with charge-dependent electron-transfer dissociation (ETD) parameters and 27% supplemental NCE. MS2 spectra for both fragmentation modes were acquired over m/z 120-2000 at 30,000 resolution, with an AGC target of 100% and maximum injection time of 250 ms.

#### LC-MS Analysis of Fabs and Reduced Chains

Antibody Fab fragments and their reduced chains were analyzed by reversed-phase liquid chromatography-mass spectrometry (LC-MS) using a Vanquish Neo UHPLC system coupled to an Orbitrap Eclipse Tribrid mass spectrometer (both Thermo Fisher Scientific), with protocols adapted from Bondt et al.[4]. Separation was performed on a MAbPac Capillary Reversed-Phase column (150 μm × 150 mm; Thermo Fisher Scientific) over a 50-min gradient at 80 °C and a flow rate of 1 μL/min. Mobile phase A was acidified MilliQ water, and mobile phase B was acidified acetonitrile. For chains, reduced using TCEP, the gradient consisted of 3 min at 10% B, a 1-min ramp to 24% B, a 50-min ramp to 36% B, a 1-min ramp to 95% B, and 2 min at 95% B. For intact Fabs, the primary gradient was a 50-min ramp from 27% to 35% B. Electrospray ionization was achieved using a Nanospray Flex Ion Source with a stainless steel emitter at 2 kV. The instrument operated in Intact Protein and Low Pressure modes. For repertoire profiling, only MS1 scans were acquired. Separate runs for Fab-level and reduced chain-level analyses included both MS1 and ETD MS2 scans. MS parameters were as follows:

| Parameter | Fab (MS1) | Fab (MS2) | Chains (MS1) | Chains (MS2) |
|---|---|---|---|---|
| Scan range (m/z) | 850-3200 | 400-3500 | 600-2400 | 400-2400 |
| Resolution | 7,500 | 120,000 | 7,500 | 120,000 |
| Normalized AGC target (%) | 250 | 1,000 | 250 | 1,000 |
| Max injection time (ms) | 50 | 246 | 50 | 500 |
| Isolation window (m/z) | - | 1.6 | - | 1.6 |
| Microscans | 5 | 10 | 5 | 10 |
| ETD reaction time (ms) | - | 5 | - | 5 |

#### Antibody Repertoire Profiling

MS1 data from LC-MS analyses of Fab fragments and reduced chains were processed and analyzed using workflows adapted from Bondt et al.[4]. Briefly, raw data were processed using BioPharma Finder version 5.1 (Thermo Fisher Scientific) with sliding window deconvolution via the ReSpect algorithm, followed by analysis in Python 3.13.7 Jupyter notebooks. Unique antibody or chain molecules (i.e., clones) were identified by their distinct combination of intact deconvoluted mass and chromatographic retention time. Absolute abundances were quantified by normalizing intensities against spiked-in internal standard mAbs. Clones were matched across samples using average linkage hierarchical clustering (unweighted pair group method with arithmetic mean, UPGMA) based on L∞ distance, with constraints on mass and retention time variations. This generated profiling datasets comprising unique molecular identifications (antibodies and chains) and their abundances in each sample, enabling longitudinal tracking of clone dynamics over time and discernment of reactivity profiles by comparing enrichments with different baits (e.g., clones captured by one SARS-CoV-2 S protein variant but not another, indicating variant-specific binding). For reactivity screening, hierarchical clustering was performed using complete linkage with correlation distance metric. Relative abundance values were $\log_{10}$-transformed prior to analysis. Missing values were imputed with a value of -1. Final visualizations of profiling data were created using the plotly and matplotlib packages in Python.

#### De Novo Peptide Sequencing and Assembly of Putative Gene Segments

Raw bottom-up LC-MS data were processed for de novo peptide sequencing using PEAKS Studio version 11.5 (build 20230821; Bioinformatics Solutions), following methods similar to Schulte et al.[5]. Precursor and fragment ion tolerances were set to 20 ppm for MS1 and 0.02 Da for MS2. Fixed modifications included carboxymethylation of cysteine (+58 Da), while variable modifications encompassed carboxymethylation of N-termini and lysine (+58 Da), oxidation of methionine and tryptophan (+16 Da), and pyroglutamic acid formation from N-terminal glutamic acid or glutamine (-18 Da or -17 Da, respectively). De novo sequenced peptides were exported as CSV files. These de novo peptide sequences were then assembled using Stitch version 1.5.0[5] (https://github.com/snijderlab/stitch) with the human germline antibody database from IMGT[6] (provided through Stitch) as templates, to generate a library of putatively assembled gene segments present in the samples. This provided an overview of germline gene segments refined by the de novo peptide data, yielding putative gene segments as they occur in the sample. These segments were subsequently connected and refined using orthogonal data, as described below.

#### Directed Serological Antibody Sequence Assembly

Integrative antibody sequence assembly was performed building on methods from Bondt et al.[4] and Schulte et al.[5]. To extend these methods to complex polyclonal samples, we combined multi-protease de novo peptide bottom-up sequencing data with ETD MS2 data of released chains and intact Fab molecules. In brief, candidate chains were matched to their parent Fabs based on orthogonal LC-MS attributes and validated manually, yielding per-antibody multi-layer data bundles as the starting point for directed sequence assembly. For each antibody, germline constant (C) and variable (V) gene segment sequences were matched to the MS2 data of its cognate released chains to identify the appropriate templates. Preliminary assembly constructs of these V-genes were then obtained from Stitch and compared to the MS2 data using Annotator[7] (https://github.com/snijderlab/annotator/). Putative mutations relative to the germlines were iteratively processed and incorporated to optimize matching with the observed MS2 evidence. Next, Fab ETD MS2 data were used alongside the chain data to match and refine the CDR3 and joining (J) gene segment of each chain. This iterative approach was combined with manual confirmation using MS2 spectra from peptides, chains, and Fabs until optimal agreement was found, assembling full serological sequences. Leucine/isoleucine residues were discerned using characteristic *w*-ions present in peptide EThcD data when available[8].

#### Genetic Analyses of Serological Antibody and BCR Repertoires

BCR sequences from the same donor (COSCA3) were obtained from the study by Brouwer et al.[1]. This dataset includes 105 unique VH and 27 unique VL sequences, of which, at that time, the 10 pairs with the highest affinity for the S protein were validated as mAbs (COVA3 series). Both serological antibody and BCR datasets were searched using IgBLAST (version 1.22.0) against V(D)J-recombined germline databases derived from IMGT[6], human entries in the CoV-AbDab[9] (excluding COVA3 sequences), and the peripheral BCR sequences themselves. For each query, the closest match in each database was selected by alignment score. Per-query and per-region (FR1, CDR1, FR2, CDR2, FR3, CDR3, FR4) mutation (deviation for CoV-AbDab) rates were computed against the closest match using Align-CLI (https://github.com/snijderlab/align-cli), allowing up to two isobaric amino acid combinations to account for ambiguities inherent to mass spectrometry-derived sequences. Differences between serological antibodies and BCRs in CDR3 lengths and mutation rates were evaluated for statistical significance using two-sided Mann-Whitney U test implemented in SciPy. Clonal expansion across germline V genes (Figure 4a and Figure S6c) was visualized as a germline-anchored radial diagram. Each sequence was placed at a radial coordinate equal to its absolute somatic hypermutation count relative to its closest IMGT V-gene match, with sequences sharing a V gene placed in a common angular sector. In addition, an UPGMA (Unweighted Pair Group Method with Arithmetic Mean) tree was constructed (Figure S6a, b) on the combined serological antibody, peripheral BCR, and germline-match sequence set. Prior to distance calculation, each query sequence was trimmed to the length of its best matching germline entry to normalize lengths across sequence sources. Pairwise distances were computed as the number of mismatched amino-acid positions plus a length-difference penalty, and the tree was built with average-linkage hierarchical clustering in SciPy. Branch positions in the polar dendrogram reflect this UPGMA topology, while leaf extension lengths are proportional to each sequence's mutation rate relative to its closest germline. All downstream analyses and figure generation were performed in Python (Jupyter notebooks and a Streamlit dashboard).

#### Recombinant Expression of Monoclonal Antibodies

Variable region sequences of the heavy and light chains were cloned into expression vectors to produce mAbs in IgG1 format as previously described[10,11]. Briefly, free-floating HEK293F cells (Invitrogen) were cultured in 293 Freestyle Expression Medium. Cells were prepared in baffled vented flasks (Corning) and used for transfection at a density of $1.0 \times 10^6$ cells/mL. For transfection of $1.0 \times 10^9$ cells, 156 μg of heavy chain plasmid DNA and 156 μg of light chain plasmid DNA were sterile-filtered and mixed with PEI-Max (Polysciences) in a 1:3 (DNA:PEI) ratio in Opti-MEM medium (Gibco). The mixture was incubated for 30 min at room temperature before addition to the cells, followed by cultivation for 6-7 days. Transfection supernatants were harvested, centrifuged, and vacuum-filtered through 0.2 μm filters. Antibodies were purified by adding immobilized Protein G agarose (Pierce) to the supernatant and incubating overnight at 4 °C with rolling. The mixture was loaded onto centrifuge columns (Pierce) and washed twice with PBS. Bound antibodies were eluted with 0.1 M glycine (pH 2.5), neutralized with 1 M Tris (pH 8.6) in a 9:1 ratio, concentrated, and buffer-exchanged to PBS using 100 kDa cut-off spin filters (Sartorius). Final proteins were sterile-filtered through 0.22 μm filters (Costar) and quantified by UV280 absorbance using the standard IgG extinction coefficient.

#### Expression of spike proteins

SARS-CoV-2 Spike and RBD protein variants were expressed transiently in HEK293F (Invitrogen, cat no. R79009) cells maintained in Freestyle medium (Life Technologies) as previously described[1,12]. Cells were transfected at a density of $1.0 \times 10^6$ cells/mL by the addition of a mix of PEImax (1.0 μg/μL) with expression plasmids (312.5 μg/L) in a 3:1 ratio in

OptiMEM. Supernatants were harvested six days post-transfection, centrifuged for 30 min at 4000 rpm and filtered using 0.22 µm Steritop filters (Merck Millipore). Glycoproteins were purified by affinity purification using Strep-TactinXT Superflow high capacity 50% suspension according to the manufacturer's protocol for gravity flow (IBA Life Sciences). Biolock solution and a 10X buffer W (1 M Tris/HCl, 1.5 M NaCl, 10 mM EDTA, pH 8.0) were diluted 1:1000 and 1:10, respectively, in the filtered supernatant prior to column loading. Protein eluates were concentrated and buffer exchanged to PBS using Vivaspin filters with a 100 kDa molecular weight cutoff (GE Healthcare). Protein concentrations were determined by the Nanodrop method using the proteins' peptidic molecular weight and extinction coefficient as determined by the online ExPASy software (ProtParam).

#### Luminex Binding Assays

Binding of serological mAbs to SARS-CoV-2 antigens was assessed using Luminex technology, following protocols adapted from Grobben et al.[13]. SARS-CoV-2 spike and RBD proteins were covalently coupled to Luminex MagPlex beads using a two-step carbodiimide reaction at a ratio of 75 µg protein per 12.5 million beads for SARS-CoV-2 Spike trimers, with other proteins (e.g., RBD) coupled equimolar to Spike trimers. Beads were washed with 100 mM monobasic sodium phosphate (pH 6.2), activated with sulfo-N-hydroxysulfosuccinimide and 1-ethyl-3-(3-dimethylaminopropyl) carbodiimide (both Thermo Fisher Scientific), and incubated for 30 min at room temperature on a rotator. Activated beads were washed three times with 50 mM MES (pH 5.0), mixed with proteins diluted in the same buffer, and incubated for 3 h at room temperature on a rotator. Beads were then washed with PBS, blocked with PBS containing 2% BSA, 3% fetal calf serum, and 0.02% Tween-20 for 30 min, washed again, and stored in PBS with 0.05% sodium azide at 4 °C. His-tag detection confirmed protein coupling levels. For assays, 50 µL of bead mixture (20 beads/µL per antigen) was incubated overnight at 4 °C with 50 µL of diluted mAb on a plate shaker. Plates were washed with TBS + 0.05% Tween-20 (TBST) using a magnetic separator, resuspended in 50 µL goat-anti-human IgG-PE (Southern Biotech), and incubated for 2 h at room temperature on a shaker. Beads were washed with TBST, resuspended in 70 µL MAGPIX Drive Fluid (Luminex), and read on a MAGPIX instrument. Median fluorescence intensity (MFI) values (from ~50 beads/well) were corrected by subtracting buffer/bead-only controls. A convalescent COVID-19 patient serum titration, positive/negative controls, and technical replicates were included for validation.

#### Pseudovirus Neutralization Assays

Neutralization activity of mAbs was evaluated using a SARS-CoV-2 pseudovirus assay adapted from Van Straten et al.[14] and Guerra et al.[15]. Pseudoviruses were generated by co-transfecting HEK293T cells with a plasmid encoding the SARS-CoV-2 S protein (WT strain with D614G mutation, Delta, and Omicron) and the pHIV-1NL43 ΔEnv-NanoLuc reporter virus plasmid (HIV-1 genome with luciferase reporter and frameshifts in env and vpr). Transfections used PEI in Opti-MEM, with supernatants harvested at 48 h, filtered (0.22 µm), and stored at -80 °C. For neutralization, serial dilutions of heat-inactivated mAbs were incubated 1:1 with pseudovirus (normalized to $10^4$-$10^5$ RLU) for 1 h at 37 °C in DMEM medium, supplemented with 10% fetal calf serum, a mixture of penicillin/streptomycin (100 U/mL and 100 µg/mL, respectively) and 1× glutamax. Mixtures were added to HEK293T/ACE2 cells (seeded at 2.0 x $10^4$ cells/well in 96-well plates 24 h prior) and incubated for 48 h at 37 °C. Culture media was removed, cells were lysed and transferred to half-area 96-wells white microplates (Greiner Bio-One). Luciferase activity of cell lysate was measured using the Nano-Glo Luciferase Assay System (Promega) with a Glomax plate reader (Turner BioSystems). Neutralization was calculated as the reduction in relative luminescence units (RLU) relative to virus-only controls. IC50 values were determined from dose-response curves fitted using nonlinear regression in GraphPad Prism. Assays included no-antibody and known neutralizing controls, with technical duplicates.

### Supplementary Figures

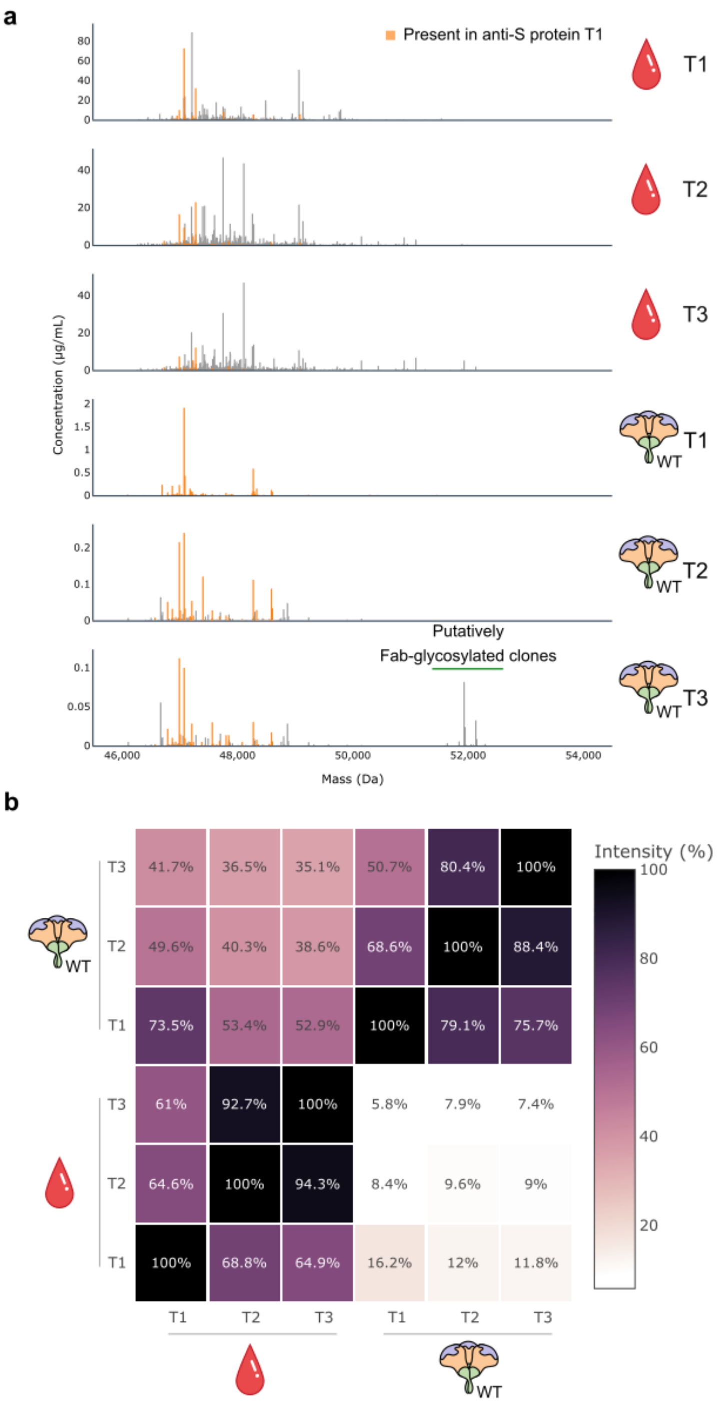

**Figure S1 | Longitudinal profiling of plasma and anti-S protein IgG1 repertoires of donor COSCA3. a,** Mass profiles showing mass and abundance of unique IgG1 Fab molecules (i.e., antibody clones) detected in plasma and derived WT S trimer enriched fractions (T1: 23 days, T2: 101 days, and T3: 158 days post symptom onset). Each stick represents a unique clone, here defined as a Fab molecule with a unique mass and retention time combination. Clones are colored based on detection in the WT S trimer enrichment from the T1 sample. Note the differences in the y-axis scales **b,** Heatmap of overlapping clonal abundance calculated from the matching clones in (a). Each row shows how much intensity in this particular sample is explained by IgG1 clones shared with other samples (columns).

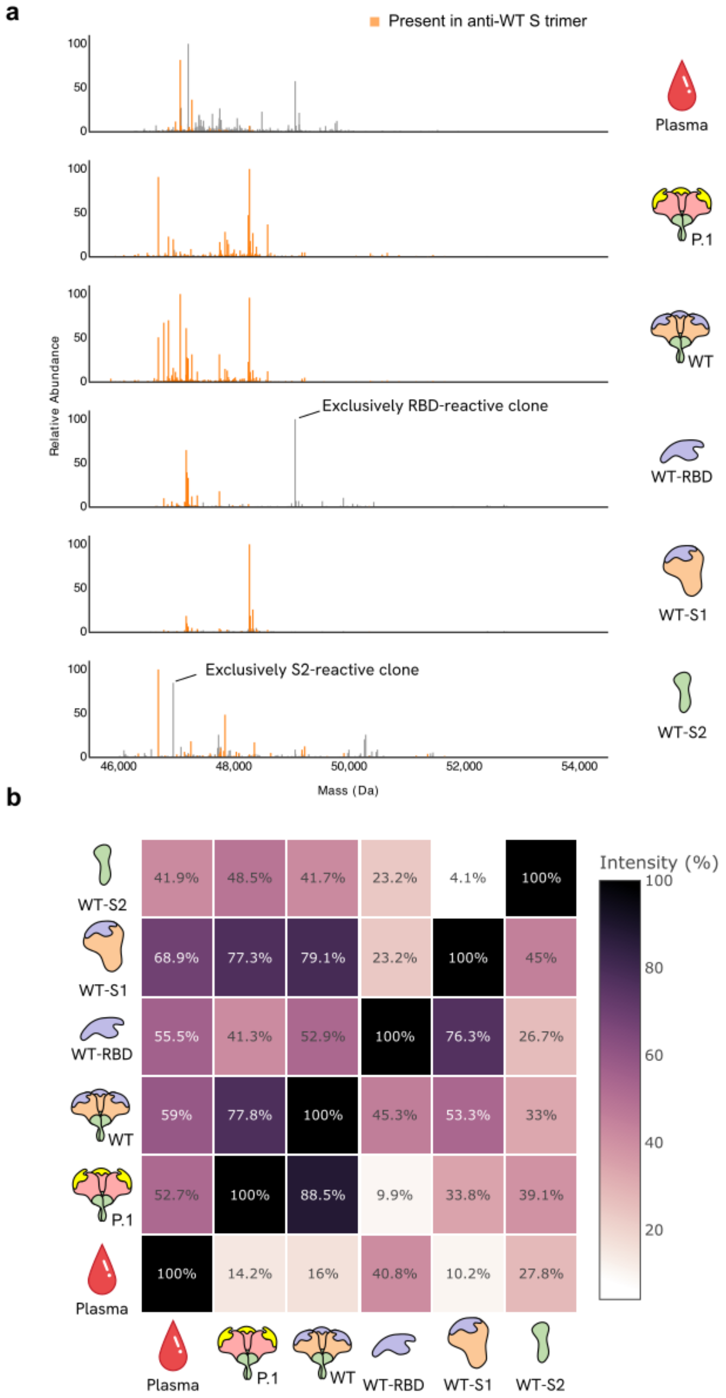

**Figure S2 | Upfront screening of IgG1 repertoires against SARS-CoV-2 S protein antigens from COSCA3 T1 plasma (23 days post symptom onset). a,** Mass profiles showing IgG1 repertoires detected in plasma and affinity enrichments using the P.1 and WT S trimers and isolated WT S1, S2, and RBD subunits. Clones are colored orange when also detected in the WT S trimer enrichment. **b,** Heatmap of overlapping clonal abundance between the detected plasma and anti-S protein IgG1 repertoires from affinity enrichments with the P.1 and WT S trimers, as well as isolated WT S1, S2, and RBD subunits.

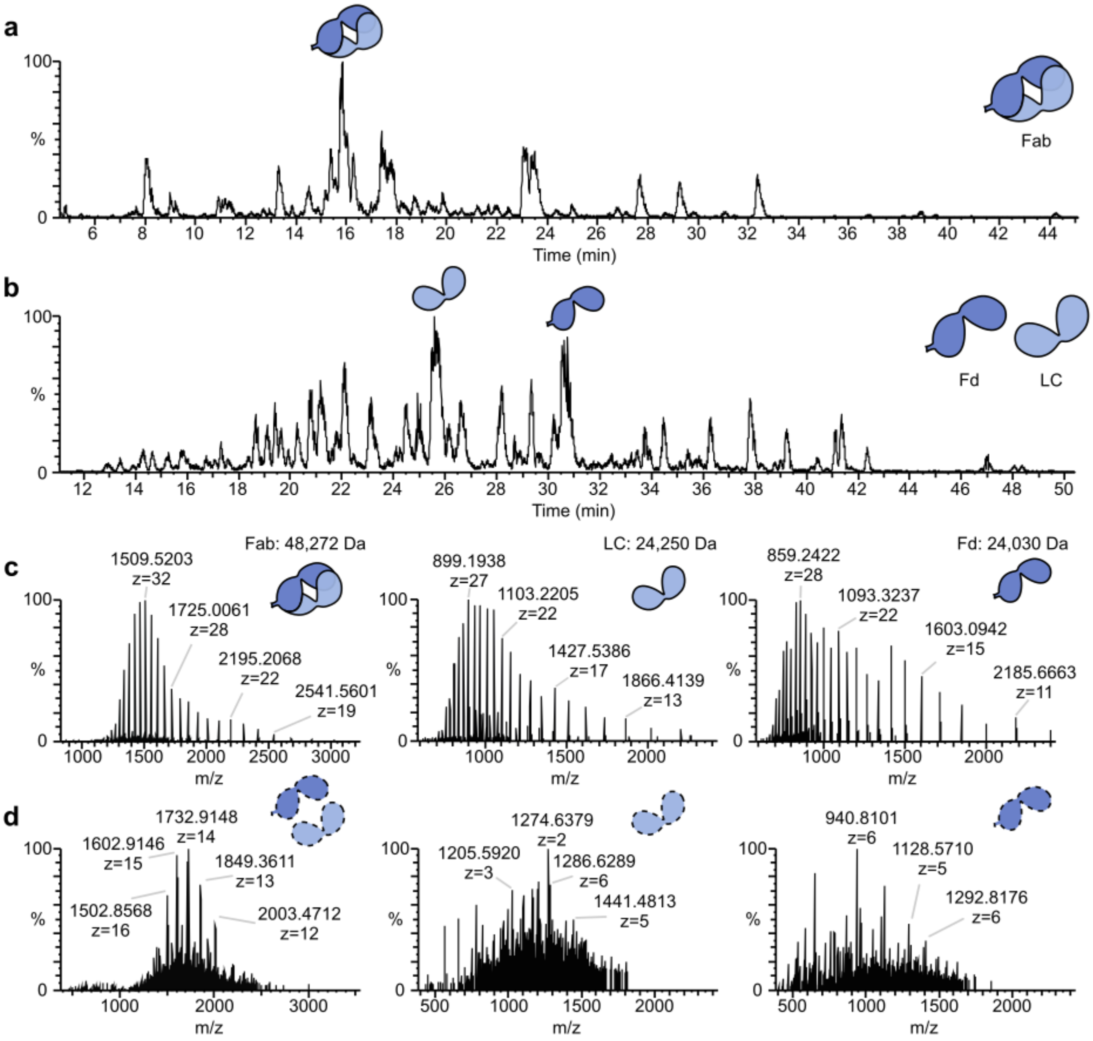

**Figure S3 | Representative LC-MS data of IgG1 Fab molecules and cognate released chains for upfront repertoire screening and integrative *de novo* sequencing of anti-S trimer clones enriched from COSCA3 T1 plasma. a,** Base peak chromatogram (MS1) of anti-P.1 S trimer IgG1 Fab molecules, showing the complexity of the polyclonal repertoire. The most abundant anti-P.1 S trimer clone (MW=47,272 Da, RT=15.8) is highlighted. **b,** Base peak chromatogram (MS1) of cognate released chains (LC and Fd) obtained through disulfide reduction of the IgG1 Fab molecules from (A). **c,** Representative MS1 spectra of an intact Fab (left), LC (center) and Fd (right) molecules of the most abundant anti-P.1 S trimer clone, providing intact mass evidence. **d,** ETD MS2 spectra of abundant charge states of the Fab (left, z=34+), LC (center, z=26+), and Fd (right, z=27+) molecules from **c**, providing fragmentation data to support *de novo* sequencing.

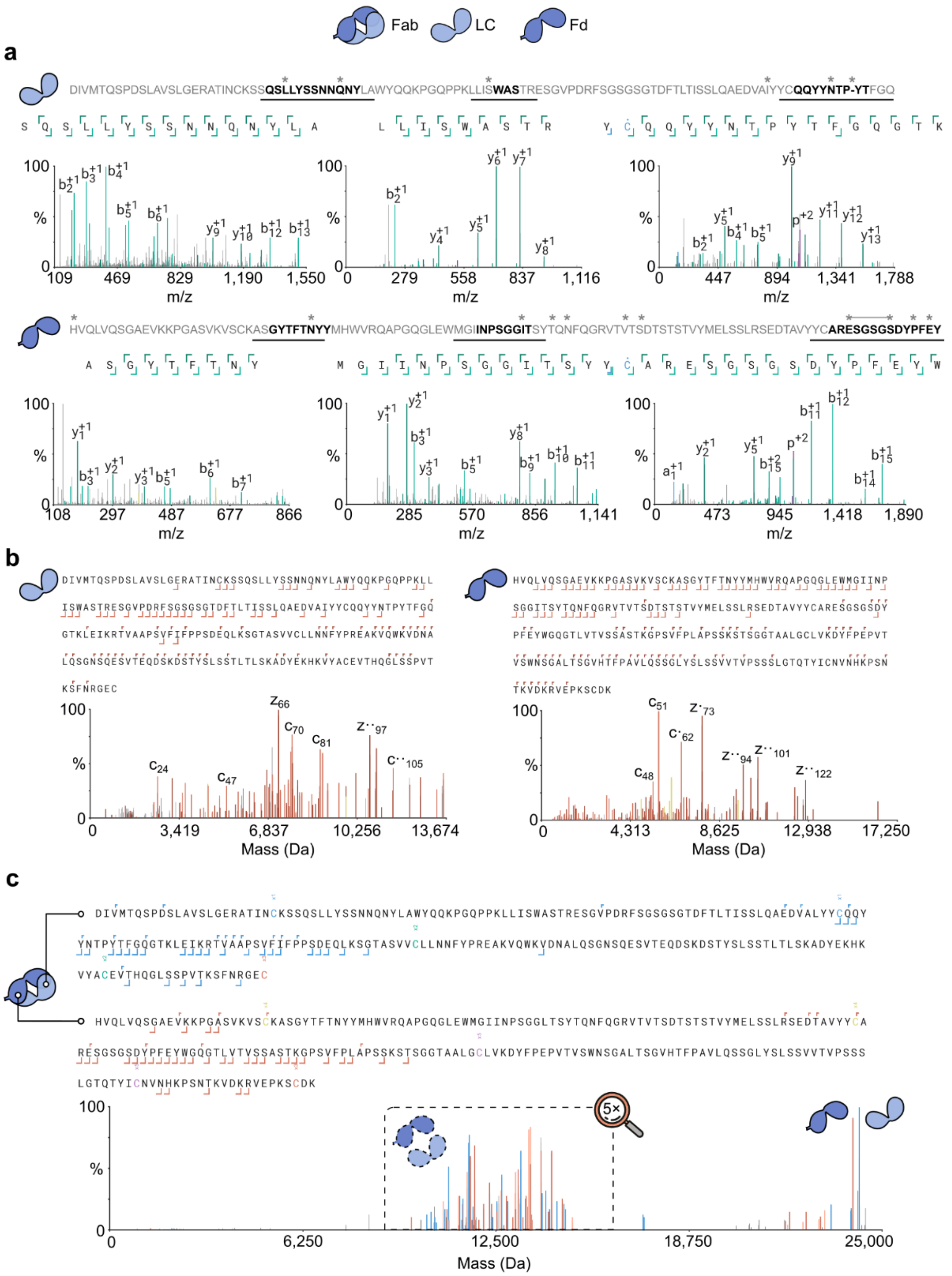

**Figure S4 | Representative peptide-level, chain-level, and Fab-level proteomics evidence for integrative de novo sequencing of anti-S trimer clones enriched from COSCA3 T1 plasma. a,** Annotated light and heavy chain variable region sequences of clone G1-CR-1 with CDRs highlighted in bold and fragmentation MS2 spectra of supporting peptides depicted below. Mutations from the closest matching germline sequences (IGKV4-1 (LC) and IGHV1-46 (HC)) are indicated with an asterisk (*). **b,** Chain-level ETD MS2 spectra of clone G1-CR-1 (LC and Fd) showing high sequence coverage and agreement with discovered sequences at a 20 ppm fragment matching tolerance. **c,** Fab-level ETD MS2 spectra of clone G1-CR-1 reveal individual masses of the constituent LC and Fd, as well as backbone fragments of CDRH3 and FR4 of both chains, matched to discovered sequences with 100 ppm fragment matching tolerance.

○ Cross-reactive ○ WT-specific

| Isotype | Clone # | Clone ID | Fab ID | Germline |
|---|---|---|---|---|
| IgG1 | 1 | G1-CR-1 | 48272 – 15.8 | IGKV4-1_IGHV1-46 |
| | 2 | G1-CR-2 | 46691 – 17.4 | IGLV3-1_IGHV3-48 |
| | 3 | G1-CR-3 | 48253 – 23.5 | IGLV2-14_IGHV5-51 |
| | 4 | G1-CR-4 | 48588 – 23.1 | IGKV4-1_IGHV5-51 |
| | 5 | G1-CR-5 | 47849 – 8.1 | IGKV1-8_IGHV3-30-5 |
| | 6 | G1-CR-6 | 48332 – 17.7 | IGKV4-1_IGHV1-46 |
| | 7 | G1-CR-7 | 46867 – 32.4 | IGLV1-40_IGHV4-39 |
| | 8 | G1-CR-8 | 48286 – 16.6 | IGKV4-1_IGHV1-46 |
| | 9 | G1-WT-1 | 47072 – 16.4 | IGLV3-25_IGHV3-23 |
| | 10 | G1-WT-4 | 46787 – 15.3 | IGLV10-54_IGHV4-39 |
| | 11 | G1-WT-5 | 47175 – 20.4 | IGKV1-9_IGHV3-66 |
| | 12 | G1-WT-7 | 47751 – 14.0 | IGKV3-15_IGHV3-21 |
| | 13 | G1-WT-9 | 47187 – 21.7 | IGKV1-9_IGHV3-66 |
| | 14 | G1-WT-10 | 47209 – 20.7 | IGKV1-9_IGHV3-66 |
| IgA1 | 15 | A1-CR-1 | 47842 – 29.2 | IGKV1-39_IGHV2-5 |
| | 16 | A1-CR-2 | 47191 – 20.2 | IGLV3-25_IGHV3-23 |
| | 17 | A1-WT-1 | 46945 – 20.8 | IGLV10-54_IGHV4-39 |
| | 18 | A1-WT-2 | 47012 – 21.7 | IGLV10-54_IGHV4-39 |

**Figure S5 | Integrative proteomics enables the discovery of 14 IgG1 and 4 IgA1 serological sequences against the SARS-CoV-2 S protein from COSCA3 T1 plasma.** Table of 18 identified and fully sequenced S protein-reactive clones: 14 IgG1 and 4 IgA1, with corresponding Fab IDs in the format [Average mass (Da) – Retention time (min)] and closest germline match.

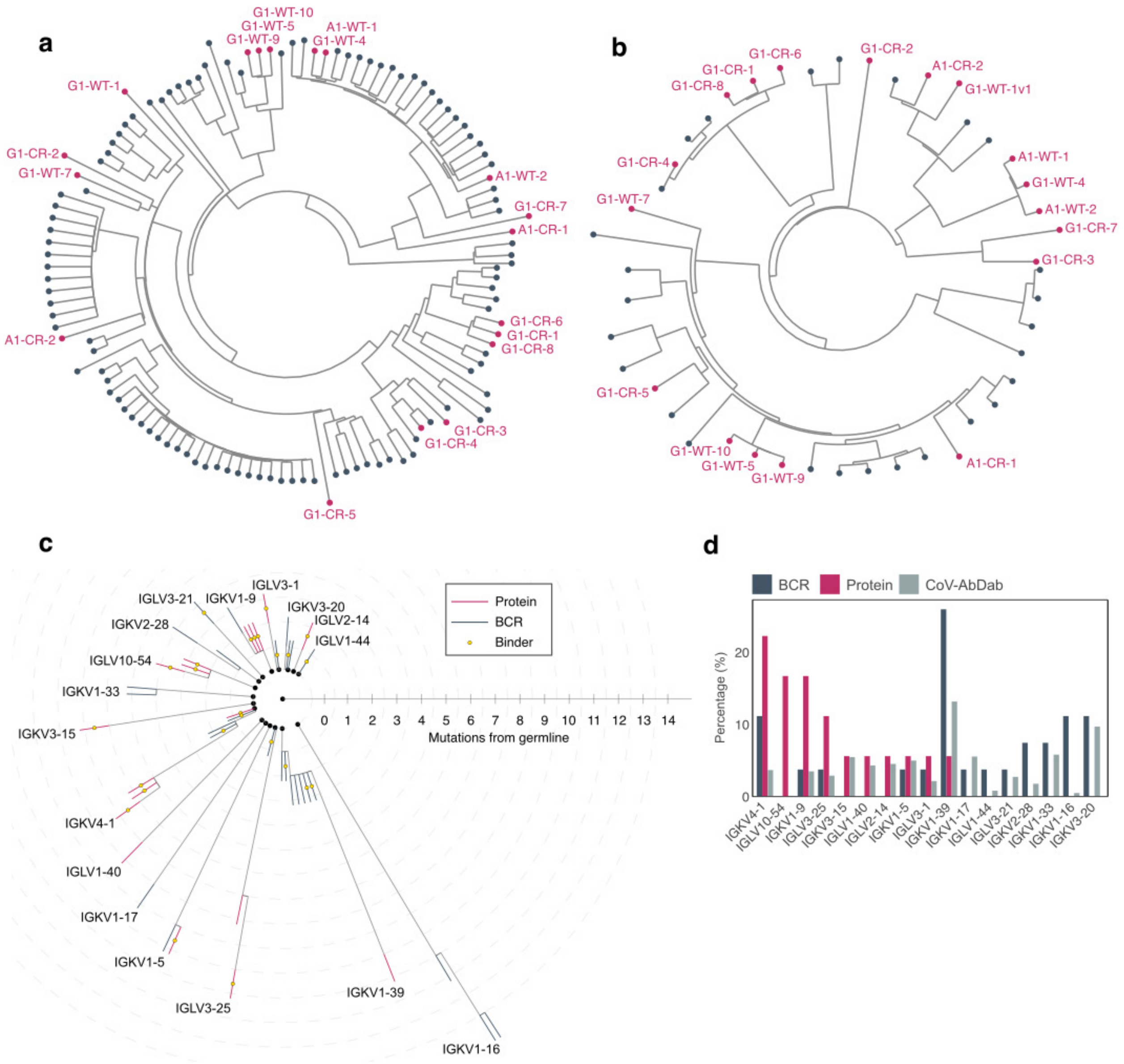

**Figure S6 | Distinctive genotypic signatures of serological antibodies and BCRs against the SARS-CoV-2 S protein. a, b,** Circular UPGMA dendrogram of heavy chain **(a)** and light chain **(b)** variable-region sequences, built from pairwise mass similarity distances against the IMGT-VDJ recombined germline database. Each leaf node represents a single sequence; dark blue nodes indicate BCR-derived sequences and pink nodes indicate serological antibody (protein)-derived sequences. Branch lengths are proportional to mass similarity distance from the germline match. Protein-derived sequences are annotated with short identifiers (e.g., G1-WT-1, G1-CR-2, A1-WT-1); prefix G1/A1 denotes antibody isotype (IgG1 or IgA1), WT = wild-type-specific clones, CR = cross-reactive clones. **c,** Germline-centered cladogram showing somatic hypermutation-based clonal relationships. Each concentric ring represents one amino acid substitution from the germline V-gene (center). Colored branches indicate query sequence types: steel = BCR-derived, crimson = protein-derived. Arc segments connect clones at identical mutation levels within germline families (angular wedges). Gold circles mark sequences present in binder-confirmed datasets. Germline families are labeled at the most mutated clone. **d,** Distribution of VL germline usage for serological antibody (red), BCR (blue), and CoV-AbDab (gray) sequences.

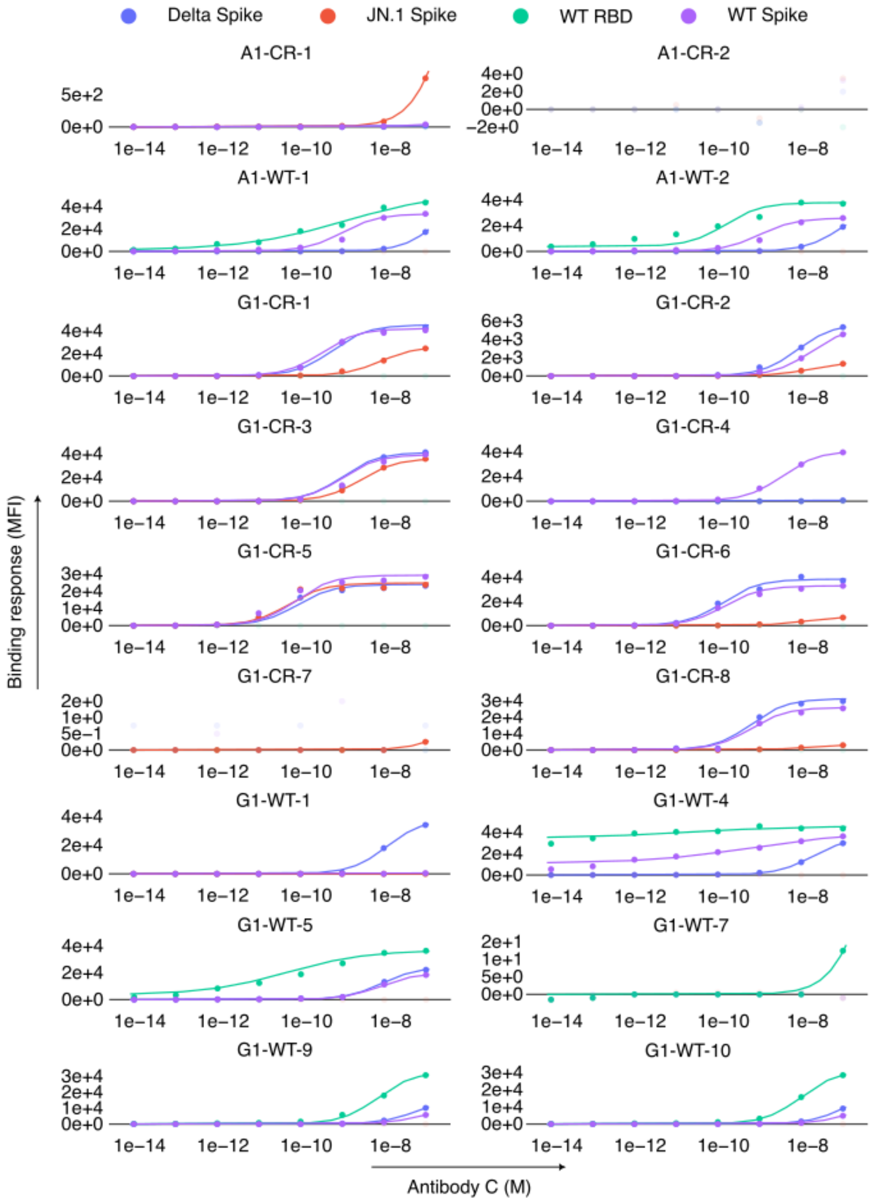

**Figure S7 | Luminex binding data for discovered anti-S protein antibodies against S protein trimer from Delta, JN.1, and WT SARS-CoV-2, as well as WT RBD.** Binding was measured in the range from 7 fM to 70 nM of antibody concentrations. Curves were fit using the Michaelis-Menten (M-M) equation when binding followed a sigmoidal curve; in cases where the M-M fit resulted in R^2 < 0.9, 4-parameter logistic regression was used for fitting. $EC_{50}$ was calculated as the concentration at half-maximum of the highest signal.
